# Antagonistic and auxin-dependent phosphoregulation of columella PIN proteins controls lateral root gravitropic setpoint angle in Arabidopsis

**DOI:** 10.1101/594838

**Authors:** Suruchi Roychoudhry, Katelyn Sageman-Furnas, Chris Wolverton, Heather L. Goodman, Peter Grones, Jack Mullen, Roger Hangarter, Jiří Friml, Stefan Kepinski

## Abstract

Lateral roots of many species are maintained at non-vertical angles with respect to gravity. These gravitropic setpoint angles (GSAs) are intriguing because their maintenance requires that roots are able to effect gravitropic response both with and against the gravity vector. Here we have used the Arabidopsis lateral root in order to investigate the molecular basis of the maintenance of non-vertical GSAs. We show that gravitropism in the lateral root is angle-dependent and that both upward and downward graviresponse requires auxin transport and the generation of auxin asymmetries consistent with the Cholodny-Went model. We show that the symmetry in auxin distribution in lateral roots growing at GSA can be traced back to a net, balanced polarization of PIN3 and PIN7 auxin transporters in the columella cells. Further, upward and downward graviresponse in lateral roots correlates with corresponding changes in PIN3 and PIN7 polar localisation. Finally, we show that auxin, in addition to driving tropic growth in the lateral root, acts within the columella to regulate GSA via the PIN phosphatase subunit RCN1 in a PIN3-dependent and PIN7-independent manner. Together, these findings provide a molecular framework for understanding gravity-dependent nonvertical growth in Arabidopsis lateral roots.

## Introduction

Gravity is one of the most fundamental, and certainly most constant, environmental signals controlling plant development. The capacity for gravity-directed growth, known as gravitropism, ensures that shoots typically grow upwards and roots grow downwards, allowing light interception and gas exchange above ground and water and nutrient uptake below. These basic processes of resource capture are enhanced enormously by the production of lateral root and shoot branches that grow out from the main root-shoot axis at non-vertical angles. Importantly, these branches are often maintained at specific angles with respect to gravity, independently from the main or parent axis from which they originate. These patterns of growth in primary and lateral organs are most easily understood in the context of the gravitropic setpoint angle (GSA) concept proposed by Digby and Firn (Digby & Firn, 1995). The GSA is the angle at which an organ is maintained with respect to gravity by the action of gravitropism. Vertically-growing organs have a GSA of 0° if they are growing towards the centre of the Earth and 180° if growing away, with non-vertical branches having GSAs between these two extremes.

To be able to alter growth according to gravity, plant organs must have the capacity to perceive their orientation within the gravity field and be able to differentially regulate elongation on the upper and lower sides of the organ. These interconnected processes are well described by two long-standing models in gravitropism research, the starch-statolith model of graviperception and the Cholodny-Went model of tropic growth. According to the starch-statolith model, orientation in the gravity field is perceived via the sedimentation of dense, starch-rich statoliths within specialised gravity-sensing statocyte cells (Nĕmec, 1900, 1901; Haberlandt, 1900; Sack & Kiss, 1989). Statolith sedimentation provides information on the organ’s angle with respect to gravity, which is then translated into tropic growth by the asymmetric redistribution of the hormone auxin to the lower side of the gravistimulated organ (Cholodny, 1927; Went, 1928; Friml et al., 2002; Ottenschläger et al., 2003; Blancaflor & Masson, 2003). According to the Cholodny-Went model, this lower side accumulation of auxin drives differential growth by inhibiting cell elongation in the root and promoting cell elongation in the shoot (Cholodny, 1927; Went, 1928). In the majority of vascular plants, statocytes are located in the endodermal starch sheath in the shoot, and the root columella cells in the root (reviewed in Blancaflor & Masson, 2003).

The starch-statolith and Cholodny-Went models are linked at a mechanistic level by the action of the PIN family of auxin efflux carrier proteins and in particular, PIN3 and PIN7 (Friml et al., 2002; Kleine-Vehn et al., 2010; Rakusova et al., 2011). In Arabidopsis, both PIN3 and PIN7 are expressed in the columella and crucially, their subcellular distribution is dependent upon the orientation of the root within the gravity field. When a primary root is growing vertically downward, PIN3 and PIN7 localisation is essentially apolar. Upon gravistimulation however, both PIN3 and PIN7 can become rapidly relocalised to accumulate preferentially on the lateral, lower-most face of the columella cells (Friml et al., 2002; Kleine-Vehn et al., 2010; Pernisova et al., 2016). This polarization of PIN3 and PIN7 within the statocytes appears to be an important early step in generating the asymmetry in auxin distribution between upper and lower halves of the root tip. Once auxin has been moved laterally from the columella, it is transported basipetally by PIN2 via the lateral root cap and epidermis (Sukumar et al., 2009; Glanc et al., 2018) to the elongation zone (EZ), where the regulation of cell expansion occurs (Rehman et al., 2010). Thus together, the starch-statolith and Cholodny-Went models, with the more recent addition of PIN-based auxin transport, provide a credible explanation for the maintenance of the vertical growth typically observed in the primary root and shoot.

Non-vertical GSAs present an intriguing problem that is fundamentally distinct from the maintenance of vertical growth. This is because in order to maintain a non-vertical GSA, root and shoot branches must by definition, have the capacity to reorient their growth both with and against gravity (Digby & Firn, 1995; Mullen & Hangarter, 2003; Roychoudhry et al., 2013; Roychoudhry & Kepinski, 2015). Previously, we established the existence of an auxin-transport-dependent antigravitropic offset (AGO or simply, offset) mechanism that counteracts the underlying gravitropic response in root and shoot branches such that stable growth at non-vertical GSAs can be maintained. In this model, GSAs are the product of the interaction of gravitropic and AGO components, with stable, angled growth occurring when both growth components are balanced (Roychoudhry et al., 2013). Importantly, while the magnitude of the AGO was proposed to be constant for a given GSA, the magnitude of the gravitropic response was predicted to increase with displacement from the vertical, as is the case in primary organs with vertical GSAs (Sachs, 1882; Mullen et al., 2000). In this way, the interplay of angle-dependent gravitropism and a non-angle-dependent AGO allows non-vertical GSAs to be maintained and regained following reorientation: if a lateral root, for example, is tilted above its GSA, gravitropic response relative to AGO activity would increase, bringing the lateral root back toward it’s GSA. In a lateral root shifted below its GSA and closer to the vertical, gravitropic response relative to AGO activity would decrease, driving upward growth. Using auxin treatment and mutants affected in either auxin homeostasis or response, this work also showed that auxin induced more vertical GSAs by diminishing the relative magnitude of the AGO (Roychoudhry et al., 2013).

While the AGO model initially provided only a conceptual framework for understanding GSA control, the fact that AGO activity is auxin-transport-dependent hints at a mechanism based in the Cholodny-Went model of tropic response. Consistent with this idea, mutation of the columella-expressed auxin efflux carriers PIN3, PIN4, and PIN7 has been shown to affect lateral root GSA (Ruiz Rosquete et al., 2013). Further, these PIN proteins also show specific expression patterns throughout the development of the emerged lateral root (Guyomarch et al., 2012; Ruiz Rosquete et al., 2013). In Arabidopsis, lateral roots (LRs) emerge at a near horizontal orientation (stage I/type 1) and following the development of differentiated columella and a distinct elongation zone that marks the acquisition of gravicompetence, undergo a brief period of downward growth (stage II/type 2) (Kiss et al., 2002; Mullen & Hangarter 2003; Ruiz Rosquete et al., 2013). From this point, lateral roots gain the capacity to stably maintain non-vertical GSAs and gradually transition through progressively more vertical GSA states, often attaining a near vertical orientation (stage II-V/type 3-6). Of these phases of gravity-dependent growth, stage III, sometimes referred to as the plateau phase, is particularly important in determining the extent of radial expansion of the root system (Mullen and Hangarter 2003; Ruiz Rosquete et al., 2013). PIN3:GFP expression in the columella is apparent from stage I onwards but begins to decline after stage III. In contrast, PIN4:GFP and PIN7:GFP are undetectable in emerging lateral roots (stages I and II), with PIN7 expression in the columella becoming apparent in stage III and PIN4 detected in the columella from ~stage IV onwards (Ruiz Rosquete et al., 2013). Based on these distinctive expression patterns it was suggested that non-vertical GSAs might simply be the result of reduced gravitropic competence, arising from the fact that PIN protein expression levels in the columella of the lateral root are lower than those in the primary root (Ruiz Rosquete et al., 2013). While the spatiotemporal expression pattern of the columella PINs is likely to be highly relevant to lateral root GSA regulation, a model based solely on gravitropic deficit is incompatible with the data supporting the GSA concept, most strikingly, the capacity of lateral roots to growth upwards if they are placed an angle that is more vertical than their GSA.

Here we have used molecular and genetic tools to dissect the mechanisms controlling GSA in the Arabidopsis lateral root. We show that unlike AGO activity, which is of essentially constant magnitude for a given GSA, gravitropism in the lateral root is angle-dependent. Further, we show that the polarity of AGO activity is, like that of gravitropic response, set and resettable relative to gravity. Critically, the kinetics of AGO resetting are slow relative to alterations in the polarity of gravitropic response. This combination of a slow-resetting and angle-independent AGO acting in tension with rapidly-modulated, angle-dependent gravitropic response underpins the maintenance of non-vertical GSAs. We show that both stable growth at GSA and upward and downward graviresponse in lateral roots are dependent on auxin transport and the generation of auxin symmetry and auxin asymmetry respectively. These patterns of auxin distribution in the root tip are reflected in the control of subcellular localisation of PIN3 and PIN7 in the lateral root columella, with the regulation of PIN3 polarity being dependent on the PIN protein phosphatase 2A subunit ROOTS CURL IN NPA1 (RCN1). Finally, we show that in addition to driving tropic response, auxin acts to induce more vertical growth by increasing RCN1 levels within the lateral root columella.

## Results

### Gravitropism in lateral roots is angle-dependent

A requirement of our previously proposed model of GSA control is that gravitropic response in lateral roots, as in primary roots, is angle-dependent (Roychoudhry et al., 2013). To test for angle-dependence in lateral root gravitropism we used a feedback-regulated system to constrain lateral roots at different stimulation angles with respect to the vertical (Mullen et al., 2000). For these experiments we analysed stage III lateral roots, which are capable of robustly maintaining non-vertical GSAs against both upward and downward reorientation (Fig. S1 A,B) (Mullen & Hangarter, 2003; Ruiz Rosquete et al., 2013; Roychoudhry et al., 2013). We measured the rate of curvature of lateral roots constrained at 30° and 45° below, and 30°, 45°, 60°, and 90° above their GSA. Constraint at 30° above or below GSA (mean GSA = 63°, SD = 7°) elicited almost identical rates of downward and upward bending respectively (Fig. 1A), consistent with the reorientation kinetics of freely-responding reorientated stage III lateral roots (Fig. S1B). Increasing the angle of reorientation to 45° above GSA led to a more than doubling of the rate of curvature (Fig. 1A). Further increases in the angle of reorientation caused a reduction in the magnitude of the response. At first sight, the angle of reorientation required for maximal response in lateral roots appears much lower than the angle of reorientation found to produce a maximal response in primary roots, which was approximately 105° (Mullen et al., 2000). However, the mean orientation of the lateral roots prior to gravistimulation was 63° (SD = 7°) from vertical, while primary roots are approximately vertical prior to reorientation. Therefore, although the angular reorientation needed to elicit a maximum response was different for primary and lateral roots, the final orientation of the root tip relative to vertical that caused the greatest response was similar for both primary and lateral roots, ~105° and ~108° respectively.

**Figure 1.**
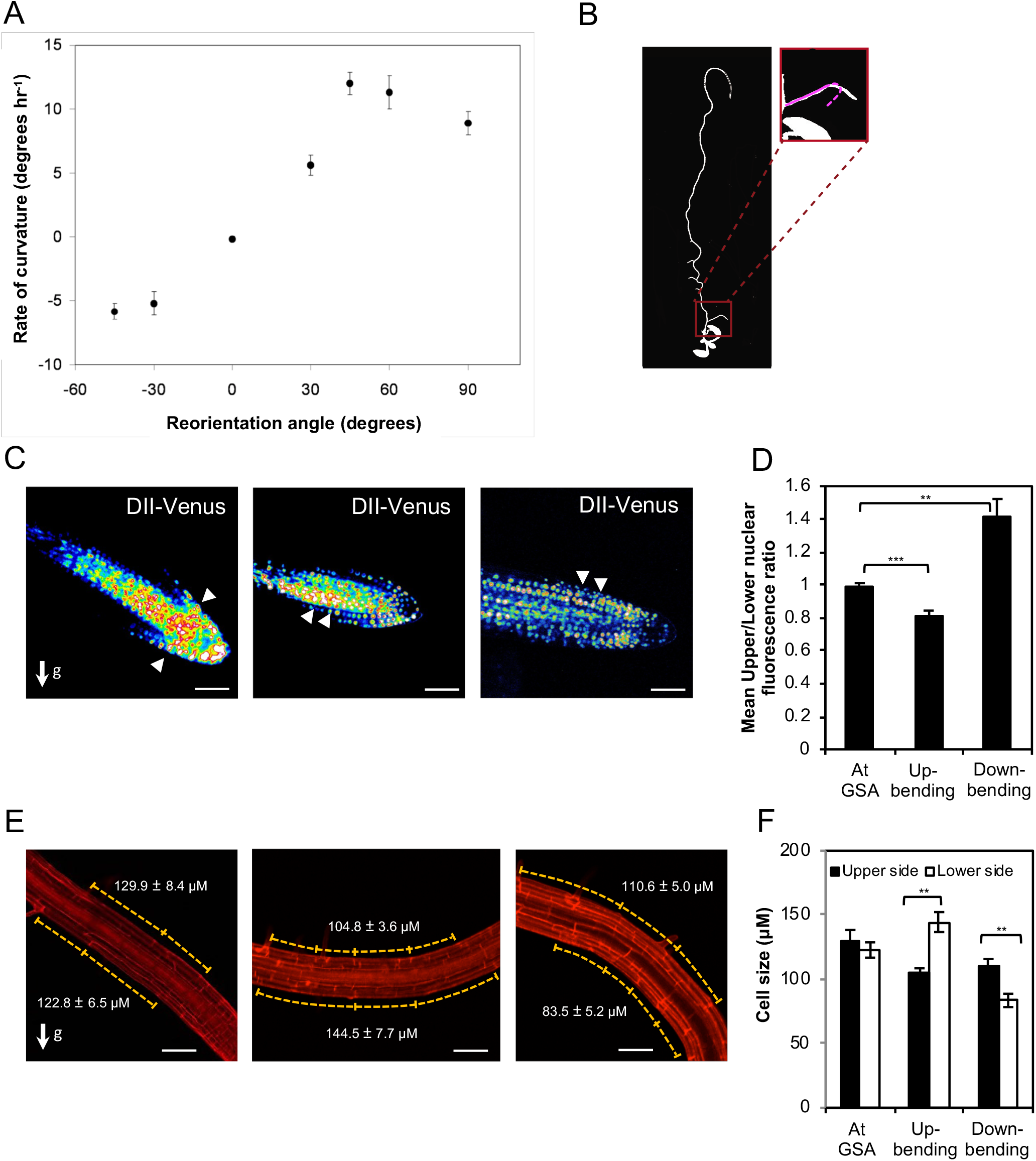
Lateral root graviresponse is angle-dependent and driven by auxin transport-dependent auxin asymmetry. (A) Dependence of gravitropic response on stimulation angle. The rate of curvature 2 hours following gravistimulation was calculated by linear regression (mean ± SE, *n* = 14-22) for lateral roots gravistimulated by varying angles of reorientation. Positive angles represent reorientations placing lateral roots above their GSA, while negative angles indicate reorientations below GSA. (B) The antigravitropic offset (AGO) is set and maintained in response to gravity. Following reorientation by 180°, a fixed, developmentally-determined AGO would result in reattainment of GSA by following a trajectory similar to that depicted by the pink dashed line in the inset image. (C) Visualization of auxin fluxes using the auxin reporter DII-Venus in upward and downward reoriented lateral roots (see Fig. S1G for quantification). Scale bar = 50 μm (D) Ratio of mean nuclear fluorescence between upper and lower halves of gravistimulated lateral roots using the auxin reporter R2D2. Note: to aid understanding, the colloquial terms up- and down-bending are used as short descriptors of lateral roots undergoing negative (upward) and positive (downward) gravitropic response respectively. (E,F) Quantification of atrichoblast epidermal cell lengths at upper and lower sides of reorientated lateral roots. In upward bending lateral roots, epidermal cells on the bottom half of the root are significantly longer than those on the upper side (E middle panel, F). In contrast, in downward bending lateral roots, epidermal cells on the bottom of the root are significantly shorter in length than those on the upper side (E third panel, F). Scale bar = 50 μm.

### The antigravitropic offset is set and maintained with reference to gravity

Our previous work has established the existence of an offset mechanism (the AGO) that counteracts positive and negative gravitropism in Arabidopsis lateral roots and shoots respectively (Roychoudhry et al., 2013). Because lateral branches in Arabidopsis and other species typically arise from a ~vertically-maintained main root-shoot axis it is possible that the offset itself could either be developmentally determined, akin to adaxial-abaxial patterning in other lateral organs, or environmentally determined, with reference to gravity. To distinguish between these possibilities, we performed a simple inversion experiment in which 12-day-old Arabidopsis seedlings were reoriented by 180° such that the previously downward faces of gravity-sensing cells would face upwards and vice versa. After 24 hours of growth we observed that lateral roots continued to maintain their original GSA but now reflected the inverted orientation of the branch relative to gravity (Fig. 1B). This finding indicates that the AGO is set, and can be reset, in reference to gravity. Were this not the case, and the offset was a gravity-independent entity, inversion would result in both gravitropism and the AGO acting across the same, new lower side of the lateral root until the root moved past the vertical, with gravitropic and AGO activity once again in opposition, resulting in a growth pattern depicted by the pink dashed line in Fig. 1B. The fact that the offset is gravity-dependent, and hence truly antigravitropic, raises the question of how AGO activity is detectable in the absence of coherent gravity stimulation under horizontal clinorotation. Under these conditions, vertically-growing primary roots and shoots without an offset will continue to grow straight as long as they are not gravistimulated prior to clinorotation. Where primary roots and shoots are gravistimulated immediately prior to clinorotation, a gravitropic bend that is directly proportional to the period of gravistimulation will continue to develop for ~1-2 hours (Hou et al., 2003, Hou et al., 2004). With this in mind, we quantified the pattern of lateral root growth under clinorotation over several hours. This showed that, despite continuing to elongate, there is no change in tip angle in the first 60 minutes of clinorotation, with the majority of bending occurring between 60 and 240 minutes (Fig. S1C). The fact that the upward bending largely ceases within 6 hours suggests that the offset is lost in the absence of unilateral gravitational stimulation, a phenomenon that is consistent with the fact that the offset is resettable if orientation with respect to gravity is changed sufficiently. Similarly, the lack of bending in the first hour of clinorotation is compatible with the idea that a lateral root growing at a non-vertical GSA can be considered to be gravistimulated. Just as in the case of gravistimulated primary roots, the effects of the resulting graviresponse are manifest during the first hour of clinorotation. The difference in clinorotated lateral roots is that this graviresponse continues to be antagonised by the activity of the offset, as it was in the upright plant. Critically, while the effects of the offset are transient in the absence of unilateral gravitational stimulation, they are longer-lived than the effects of gravitropic response. This difference in the timing of loss of gravitropic and antigravitropic growth components drives the observed upward bending under clinorotation. It follows that the patterns of growth on the clinostat and under reorientation in the upright plant are two manifestations of the same mechanism. In both cases, upward growth occurs because of a reduction in the magnitude of the gravitropic component. Under clinorotation, this reduction is the result of the withdrawal of unilateral gravitational stimulation, while in the lateral root placed more vertically than its GSA, the reduction is attributable to the angle-dependence of gravitropic response in lateral roots (Fig. 1A).

### Gravitropic auxin transport is offset by an antagonistic auxin flux in Arabidopsis lateral roots

The control of auxin distribution across the root tip is central to the maintenance of vertical GSA sin the primary root. To begin to explore the role of auxin transport in the maintenance of non-vertical GSAs, we tested the effect of the auxin transport inhibitor NPA on both upward and downward gravitropic growth in reorientated lateral roots. In these experiments, lateral roots treated with either 0.2 μM or 0.4 μM NPA failed to return to their original GSA after 24 hours following rotation by 30° either above or below their GSA (Fig. S1E,F). At both concentrations, the growth rates of lateral roots are not significantly different from the wild type in our growth conditions (Fig. S1D) (Roychoudhry et al., 2013). These data therefore indicate that auxin transport is necessary for both upward and downward gravity induced growth curvatures.

To analyse auxin distribution and response in lateral roots growing at their GSA, and following gravistimulation above and below their GSA, we used the reporters DII-Venus and the ratiometric DII-Venus variant, R2D2 (Band et al., 2012; Li et al., 2015). In lateral roots growing at their GSA, these reporters indicated no significant difference in auxin levels between the upper and lower halves of lateral roots (Fig. 1C,D; Fig. S1G). The inference of auxin levels from DII-Venus and R2D2 in this context is further supported by the lack variation in TIR1/AFB auxin receptor levels across the lateral root, quantified by TIR1:Venus, AFB2:Venus, and AFB3:Venus translational reporter expression (Fig. S1H) (Roychoudhry et al., 2017). In lateral roots reorientated upwards, above their GSA (and bending downwards), R2D2 and DII-Venus signals indicated higher levels of auxin accumulation on the lower side of lateral roots (Fig. 1C,D; Fig. S1G). Conversely, in roots displaced below their GSA (and thus bending upwards), R2D2 and DII-Venus signals indicated higher levels of auxin accumulation on the upper side of the lateral root, in the direction of tropic growth (Fig. 1C, D, Fig. S1G). Taken together, these data indicate that the maintenance of non-vertical GSAs is auxin transport-dependent and entirely consistent with the Cholodny-Went model of tropic growth.

To understand how these patterns of auxin distribution in reorientated lateral roots relate to the generation of tropic curvature we measured atrichoblast epidermal cell lengths across the upper and lower sides of stage III lateral roots, both at and displaced from their GSA. Consistent with the lack of auxin asymmetry in lateral roots growing at their GSA, we found that there were no significant differences in cell lengths between the upper and lower sides of the root (Fig. 1E,F). In downward bending lateral roots, epidermal cells on the lower side of the root were significantly shorter than those at the upper side, consistent with the asymmetric auxin accumulation in these cells and resulting auxin-mediated growth inhibition (Fendrych et al., 2018) in the lower half of the root (Fig. 1E,F). In upward bending roots, we observed that the cells on the lower side of the root were significantly longer than those on the upper side, and, interestingly, they were also longer than epidermal cells in roots growing at their GSA (Fig. 1E,F). Importantly, epidermal cells on the upper side of lateral roots undergoing either upward and downward tropic growth responses did not differ significantly in length. These data indicate that tropic growth in lateral roots is driven principally by control of cell elongation on the lower side of the root. In the context of the auxin distribution data, this would be consistent with the idea that stimulation angle-dependent variation in the transport of auxin to the lower side of the lateral root, against a more constant and angle-independent flux of auxin to the upper side, is the mechanism driving both upward and downward tropic growth.

### Polarity and distribution of PIN proteins in lateral roots

The auxin efflux carriers PIN3 and PIN7 have been previously described to play a major role in translating information on the direction of gravity into asymmetric auxin fluxes by their gravity-induced polarization (Friml et al., 2002; Kleine-Vehn et al., 2010; Rakusova et al., 2011). Both PIN3 and PIN7 are expressed in stage III lateral roots, from which point GSAs are robustly maintained (Guyomarc’h et al., 2012; Ruiz Rosquete et al., 2013). We therefore decided to study in detail the localisation and distribution of these PINs in Arabidopsis lateral roots growing at non-vertical GSAs using vertical-stage confocal microscopy (von Wangenheim et al., 2017. In agreement with previous data (Ruiz Rosquete et al., 2013; Guyomarc’h et al., 2012), we found that PIN3 was expressed mainly in the top two tiers of columella cells. Here, PIN3 largely showed a bipolar distribution with respect to the upper and lower side walls of columella cells relative to gravity (45% of roots analysed). In addition, a significant proportion of roots had polar distribution of PIN3, with approximately 21% of roots having an ‘upward’ polarity and 34% having a ‘downward’ polarity (Fig. 2A,E; Fig. S2A). In contrast, PIN7 had a distinctly different polarity distribution: PIN7 had upward polarity distribution in over 50% of lateral roots analysed, and downward and apolar distributions in approximately 26% and 22% of lateral roots respectively (Fig. 2B,E). In contrast with lateral roots at GSA, in primary roots placed non-vertically (~45°), both PIN3 and PIN7 polarised predominantly towards the lower side of the columella in agreement with previous studies (Fig. 2C,D) (Kleine-Vehn et al., 2010).

**Figure 2:**
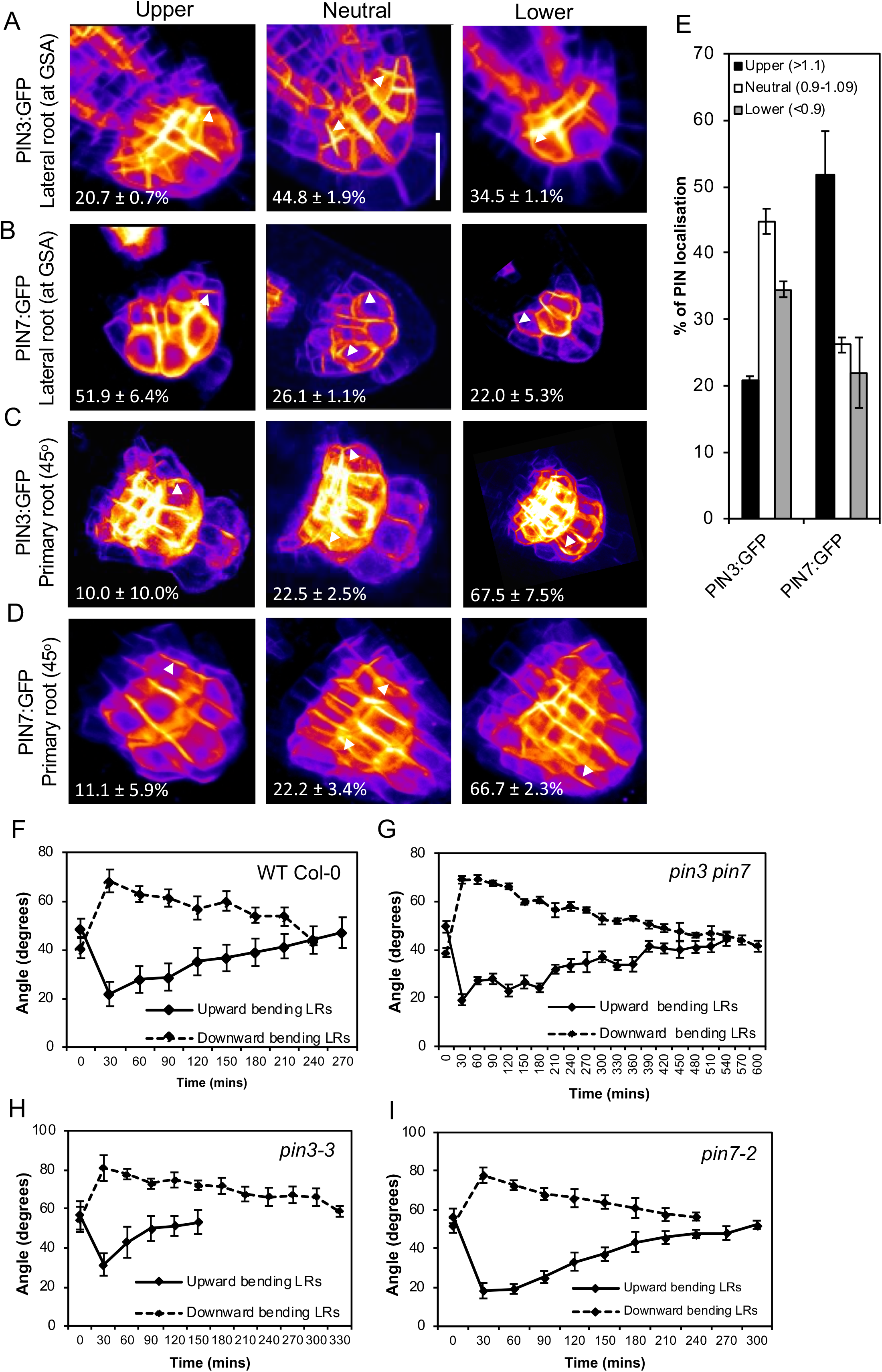
PIN polarity distribution in lateral roots. (A-E) Quantification of PIN polarity in lateral roots of seedlings expressing PIN3:GFP (A, E) and PIN7:GFP (B,E). Fluorescence was measured on upper and lower membranes of outer columella cells as indicated in Fig. S2A. PIN polarities were assigned based on ratios of upper/lower mean fluorescence values. (<0.9 = lower, 0.91 – 1.1 = neutral/dipolar, >1.1 = upper). PIN3 shows a prominent lower bias (A,E). In contrast, PIN7:GFP shows a prominent upper bias (B,E) in lateral roots growing at their GSA. However, both PIN3:GFP (C) and PIN7:GFP (D) are predominantly polarized towards the lower plasma membrane in primary roots reoriented by ~45°. Scale bar = 15 μm (C,D). Comparison of reorientation kinetics in lateral roots of 12-day-old WT Col-0 (F), *pin3 pin7* (G) *pin3-3* (H) and *pin7-2* (I) seedlings gravistimulated both above and below their GSA. Average GSA of 10-12 upward and downward bending stage III lateral roots was quantified after reorientation until the roots were within 5 degrees of their original GSA. *pin3* lateral roots reorientate upwards significantly faster (H), while *pin7* lateral roots reorientate downwards at a faster rate (I). WT Col-0 control lateral roots reorientate back to their GSA in both directions in approximately 6 hours (F). In contrast, reorientation in both directions is delayed in the *pin3 pin7* double mutant (G).

To explore the significance of these cell biological observation, we compared the kinetics of graviresponse of *pin3-3* and *pin7-2* single and double mutant lateral roots to those of wild-type Col-0. Upon reorientation by 30° above and below GSA, wild-type lateral roots returned to their original GSA in approximately 6 hours (Fig. 2F). Consistent with their apparent role in GSA control, lateral roots of the *pin3 pin7* double mutant were severely delayed in returning toward their GSA, although both upward and downward tropic growth was still apparent (Fig. 2G). Because previous studies have shown that PIN4 expression domains expand into the columella in a compensatory manner in the *pin3* and *pin7* mutant backgrounds (Vieten et al., 2005), we also examined the graviresponse in the *pin3 pin4 pin7* triple mutant and found the response to be further reduced (Fig. S1I).

The graviresponse kinetics of the *pin3* and *pin7* single mutants were particularly interesting. We found that lateral roots of *pin3* seedlings reorientated upwards significantly faster than downwards (Fig. 2H), while the reverse was true for lateral roots of *pin7* seedlings, albeit to a lesser degree (Fig. 2I). We also tested the response of the *pin3 pin4* double mutant, which, similar to the *pin3* single mutant, exhibited a much more rapid upward relative to downward tropic growth (Fig. S1J). These data indicate that the rapid upward bending associated with loss of PIN3 function requires PIN7 and are thus consistent with the observed subcellular polarity bias of PIN7 and to some extent, PIN3.

Previous studies have demonstrated that changes in PIN localisation in columella and endodermal cells following gravistimulation of primary roots and shoots leads to auxin redistribution and graviresponse (Friml et al., 2002, 2003; Kleine-Vehn et al., 2010; Rakusova et al., 2011). To establish if such changes were also apparent in lateral roots, we gravistimulated lateral roots both above and below their GSA and examined PIN3:GFP and PIN7:GFP localisation in the columella by vertical-stage confocal microscopy (von Wangenheim et al., 2017). In these experiments we measured the ratio of GFP signal on the upper and lower sides of outermost flanking cells. For lateral roots growing at their GSA, the upper/lower ratio for PIN3:GFP was approximately equal to one, whereas for PIN7:GFP the ratio was slightly higher (1.2) for PIN7:GFP (Fig. S2B-E), as described above (Fig. 2B). Stimulation above GSA (downward bending) shifted the polarity of both PIN3 and PIN7 to a predominantly basal localization, similar to that in primary roots (Fig. S2B-E). In contrast, where lateral roots were reorientated below their GSA (upward bending), we observed an increased signal at the upper plasma membrane relative to the lower side for both PIN3:GFP and PIN7:GFP lateral roots (Fig. S2B-E). Thus, these shifts in polarity of PIN3 and PIN7 are consistent with the observed R2D2 and DIIV data demonstrating auxin redistribution during lateral root gravistimulation.

In addition to PIN3 and PIN7 activity in the columella, root graviresponse also requires the action of PIN2 in the epidermis to drive the basipetal flow of auxin away from the root tip. We therefore also studied PIN2:GFP expression in lateral roots growing at their GSA to check for any differential expression of PIN2 that might contribute to the regulation of growth angle. Confirming previous studies (Lofke et al., 2015), we observed slight differential expression of PIN2:GFP between trichoblast and atrichoblast cell files (Fig. S2F) but did not observe differences in PIN2 expression between upper and lower halves of lateral roots (Fig. S2G), in contrast to those observed in gravistimulated primary roots (Baster et al., 2013).

### PIN protein retention at the plasma membrane differs between upper and lower faces of repolarising lateral root statocytes

The subcellular distribution of PIN proteins is regulated via cycles of endocytosis and polar or apolar redelivery to the plasma membrane (reviewed in Adamowski & Friml, 2015). In order to understand if there are differences in PIN stabilisation within the upper and lower side membranes of lateral root statocytes at GSA, we designed an assay to capture the dynamics of PIN protein relocalisation during the gravity-induced repolarization of the cell. This involved ‘flipping’ lateral roots growing at their GSA by 90° within their axis of growth, simply by moving from vertical-to horizontal-stage confocal microscopic imaging and analysing, over time, the faces of the statocyte that were previously ‘up’ and ‘down’ relative to gravity prior to the ‘flip’ (Fig. S3A). For both PIN3:GFP and PIN7:GFP, the ratio of fluorescence signals at the upper and lower lateral root columella cell faces was recorded immediately after flipping and then at 30 minute intervals for 2 hours. We quantified PIN polarization as the percentage of roots that had a ratio of upper-to-lower side PIN:GFP signal that was markedly different from 1 (lower than 0.9 or greater than 1.1). Consistent with previous experiments (Fig. 2A,B,E), we found that immediately following the flip, PIN3:GFP showed a symmetrical to lower side bias (Fig. 3A,B), while a symmetrical to upper side bias was evident for PIN7:GFP (Fig. 3C,D). Thirty minutes after ‘flipping’ we found that the majority of lateral roots now displayed increased PIN3 and PIN7 localisation towards the former upper cell side (Fig. 3A-D). This indicates that lower side PINs are endocytosed and polarised in the new direction of gravity and statolith sedimentation at a faster rate as compared to upper side PINs. Comparing the polarity distribution throughout the course of the experiments, we found that for both PIN3 and PIN7 the proportion of lateral roots with upper polarity decreased, while that of symmetrical or ‘neutral’ polarity increased over time. Notably, this progression to a neutral state of polarity occurred more slowly for PIN7 relative to PIN3 (Fig. 3B,D). For both PIN3 and PIN7, the majority of lateral roots had acquired a symmetrical distribution across both cell sides two hours after flipping (Fig. 3A-D). As a control, we performed the same assay with another plasma membrane protein marker line, WAVE:YFP (Geldner et al., 2009). In lateral roots at their GSA, there was no asymmetry in WAVE:YFP expression across the upper and lower membranes of columella cells (Fig. S2H,I). Additionally, ‘flipping’ did not lead to the generation of any asymmetry of WAVE:YFP across the cell, suggesting that the ability to repolarize in the direction of gravity is not a general property of plasma membrane proteins. Taken together, these results indicate that there is differential stability of PIN3 and PIN7 at the upper versus lower plasma membranes of lateral root statocytes growing at GSA, with PIN3 and PIN7 being retained at the upper sides for longer.

**Figure 3:**
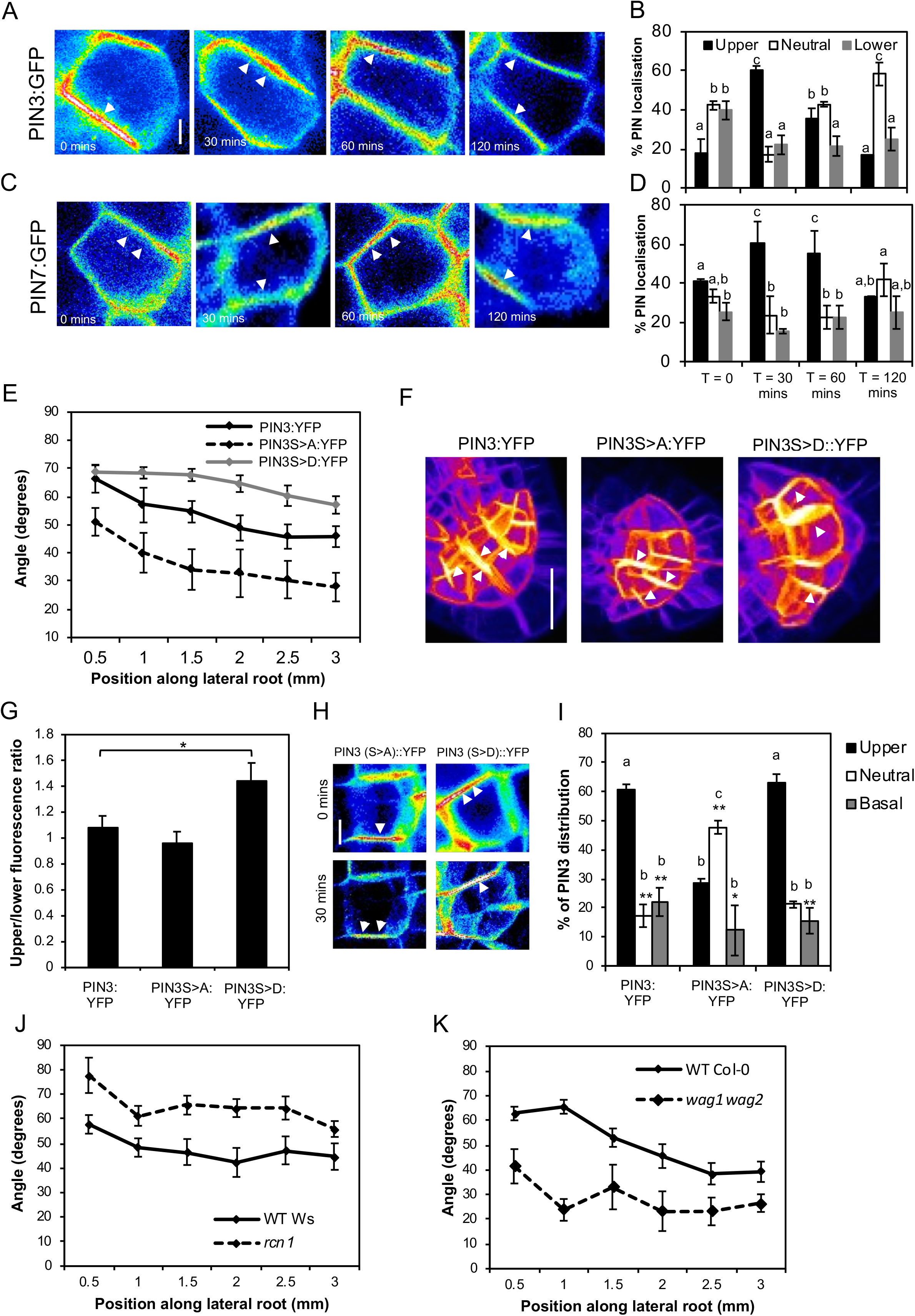
Phosphorylation increases PIN stability on the plasma membrane. (A-D) PIN polarity distribution in columella cells of lateral roots rotated around their axis of growth by 45° (‘flip assays’, see Figure S3A for a diagrammatic description of the experiment). In all panels, the former upper side of the columella cell is towards the top of the page. Post flip, phosphorylated PIN3 and PIN7 are retained on the upper plasma membrane for approximately 2 hours longer than lower side unphosphorylated PINs. PIN3 (A,B) and PIN7 (C,D) polarity gradually becomes symmetrical (neutral) on upper and lower sides of the plasma membrane 2 hours after ‘flipping’ (D). Letters (a,b,c,d) represent p values of < 0.005. (E-G) Quantification of GSA phenotypes (E) and PIN3 polarity (F,G) in transgenic lines expressing nonphosphorylatable (S>A) or phosphomimic (S>D) variant of PIN3-YFP. * represents a p value < 0.05. Scale bar = 15 μm (H,I) PIN3:YFP polarity distribution in horizontally flipped lateral roots of PIN3S>A:YFP and PIN3S>D:YFP. PIN3:YFP signal becomes equally distributed 30 mins after flipping in the PIN3S>A:YFP phosphovariant line, but remains polarized to the upper side of the cell in PIN3S>D:YFP lateral roots. Letters (a,b,c) represent p values < 0.005. (J,K) Quantification of GSA phenotypes in *rcn1* and *wag1 wag2* mutants. *rcn1* lateral roots have significantly less vertical lateral roots as compared to WT Ws seedlings (J). In contrast, *wag1 wag2* seedlings have significantly more vertical lateral roots than WT Col-0 controls (K). Scale bar = 5 μm in (A), (C) and (H).

### PIN phosphorylation affects PIN polarity and redistribution kinetics in lateral root statocytes

Previous reports have demonstrated that the polarity of PIN1 and PIN2 in epidermal and vascular cells is dependent on their phosphorylation state, mediated by the PINOID/WAG kinase- and PP2AA phosphatase families (Zhang et al., 2009; Dhonushke et al., 2010; Friml et al., 2004; Michniewicz et al., 2007). These studies showed that phosphorylation of specific serine (S) residues in the cytoplasmic loops of these membrane proteins induces localisation to the shootward (upper) plasma membrane, while their dephosphorylation causes PINs to localise to the rootward (lower) plasma membrane. To begin to explore the role of phosphorylation in the subcellular distribution of PINs within the lateral root columella we analysed lateral root GSA of PIN3:YFP phosphovariant lines in which known PID/WAG- or D6PK-targeted serine residues were mutated to either a nonphosphosphorytable alanine (A) (Grones et al., 2018; Zourelidou et al., 2014) or to the phosphomimic amino acid aspartic acid (D) (Grones et al., 2018). Previous studies have shown that phosphorylation at the specific residues of S316, S317 and S321 for PID/WAG and S215 and S283 (annotated as S4 and S5) for D6PKs can affect the gravity-induced repolarization of PIN3 in the root and hypocotyl respectively (Grones et al., 2018; Zourelidou et al., 2014). We found that lines in which the S316, S317 and S321 PID/WAG sites of PIN3:YFP were mutated to alanine (PIN3S>A:YFP) had more vertical lateral root GSAs, while the mutation of those same residues to aspartic acid (PIN3S>D:YFP) induced lateral roots to grow at a more horizontal GSA than control PIN3:GFP plants (Fig. 3E). In contrast, the mutation of D6PK phosphosites to alanine (PIN3:S4S5A:YFP) had no effect on lateral root GSA (Fig. S3B).

Consistent with their effect on lateral root GSA, we observed that, compared to native PIN3:YFP, the distribution of PIN3S>D:YFP was shifted significantly towards the upper membrane of lateral root columella cells, while that of PIN3S>A:YFP was shifted slightly, but not significantly towards the lower membrane (Fig. 3F,G). We also performed flip assays using the PIN3:YFP phosphovariant derivatives, which showed that the characteristic persistence of a stronger PIN3 signal of the former upper side of lateral root statocyte at 30 minutes post-flip was lost in the PIN3S>A:YFP line but retained in PIN3S>D:YFP (Fig. 3H,I).

The absence of a GSA phenotype in the D6PK phosphovariant line prompted us to focus on the role of the PID/WAG kinases and PP2AA/RCN1 phosphatases on the regulation of PIN-mediated transport from lateral root statocytes. Analysis of transcriptional and translational marker lines showed that while PID and WAG1 were below the level of detection in primary and lateral root columella cells, a third member of this family, WAG2 is expressed solely in the lateral root columella (Fig. S3C). RCN1 is expressed in both primary and lateral root columella cells (Fig. S3C). Interestingly, loss of RCN1 function in the *rcn1* mutant causes lateral roots to grow with a significantly less vertical GSA (Fig. 3J), while the double loss-of-function mutant *wag1 wag2* induces a more vertical lateral root GSA (Fig. 3K). These data are thus consistent with the PIN phosphovariant data and the idea that RCN1-mediated phosphatase activity facilitates the ‘downward’ fluxes of auxin from lateral root statocytes, and that kinases such as WAG2, and possibly others that target the PIN3 cytoplasmic loop, facilitate an opposite, ‘upward’ auxin flux.

### Auxin regulates lateral root GSA through a PIN3-specific phosphorylation module

It has previously been shown that auxin treatment is able to shift lateral root GSA towards a more vertical orientation (Ruiz Rosquete et al., 2013;; Roychoudhry et al., 2013; Roychoudhry et al., 2017). We therefore hypothesised that auxin might affect lateral root GSA by affecting PIN polarity, for example, by increasing the pool of dephosphorylated PINs within the lateral root columella. This increase could be achieved either through an auxin-mediated up-regulation of RCN1 expression or activity and/or down-regulation of the opposing kinase expression or activity. To explore this possibility, we tested the effect of auxin treatment on lateral root GSA in *rcn1* and *wag1 wag2. rcn1* lateral roots failed to respond to auxin treatment (Fig. 4A), while the lateral roots of *wag1 wag2* double mutants shifted to a more vertical GSA orientation, similar to wild-type (Fig. S4A,B), indicating that auxin might control lateral root GSA through an RCN1-dependent pathway. Consistent with this idea, the overexpression of *RCN1* driven specifically in the columella by the promoter of *ARL2*, a columella-specific gene (Harrison & Masson, 2008) (Fig. S4C), in the Col-0 background led to a significantly more vertical lateral root GSA phenotype (Fig. 4B). Indeed, this same *ARL2::RCN1* transgene was able to rescue the horizontal GSA phenotype of *rcn1* lateral roots (Fig. 4E).

**Figure 4:**
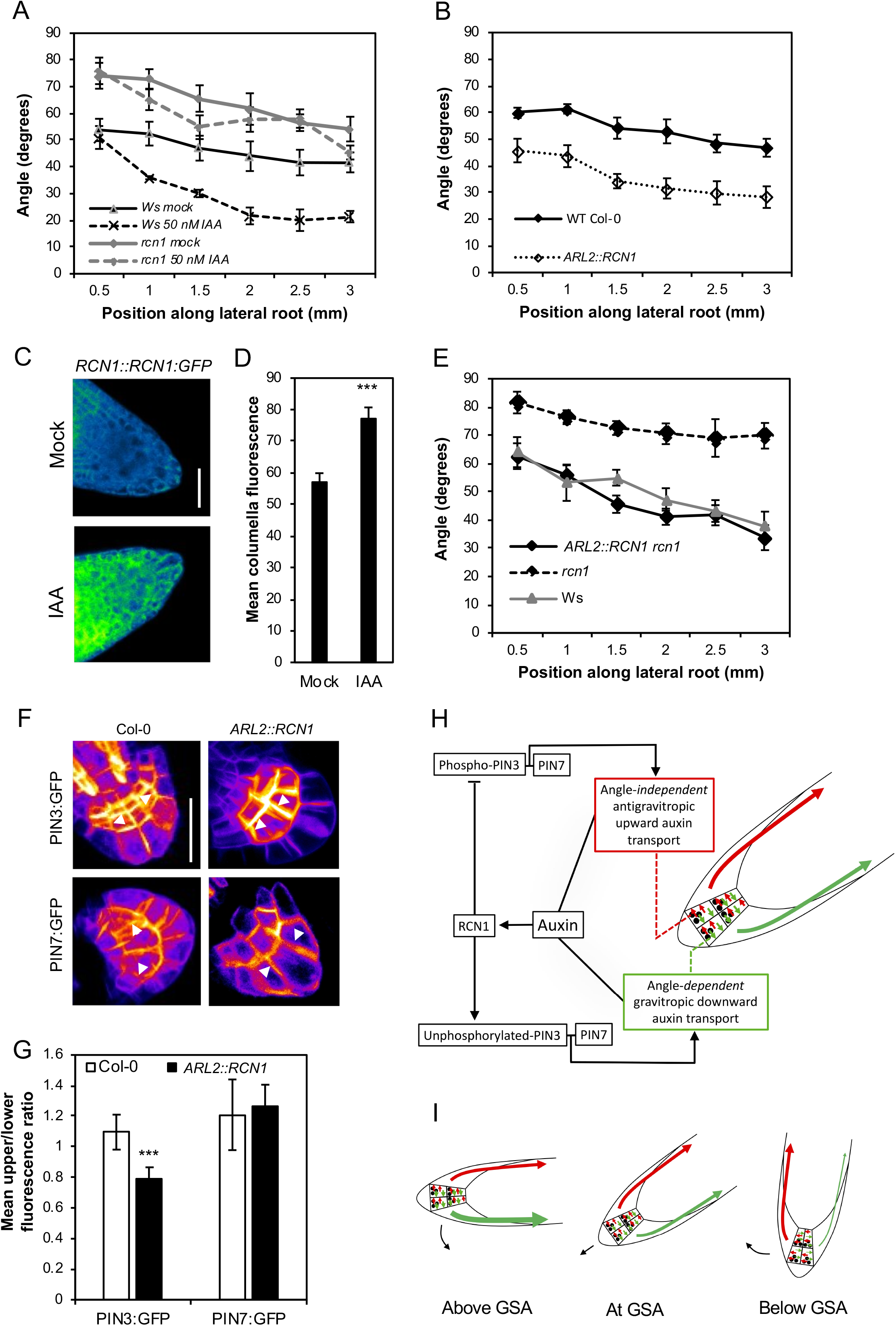
Auxin regulates lateral root GSA through an RCN1-dependent PIN3 module. *rcn1* lateral roots are insensitive to the auxin effect on GSA (B) Overexpression of *RCN1* driven by the *ARL2* promoter (ARL2::RCN1) in a WT Col-0 background results in a significantly more vertical lateral root GSA phenotype in contrast to Col-0 control. (C,D) RCN1:GFP protein levels in ten day old lateral roots treated with 50 nM IAA for 4 hours. Auxin treatment results in a significant increase (p < 0.001 (5.06661E-05) in GFP signal in the columella cells of RCN1:GFP lateral roots (D). Scale bar = 20 μm in (C). (E) Columella specific expression of *RCN1* driven by the *ARL2* promoter is sufficient to rescue the horizontal GSA phenotype of *rcn1* lateral roots. (F,G) Overexpression of *RCN1* in lateral root columella cells leads to a significant shift in PIN3:GFP polarity towards the lower side of the cell. In contrast, PIN7:GFP polarity is unaffected. ** represent a p value of <0.001, scale bar = 15 μm. (H) Model of GSA control in the lateral root in which phosphorylated PIN3 & PIN7 mediate upward, antigravitropic auxin flux from columella cells, while unphosphorylated PIN3 & PIN7 mediate downward, gravitropic auxin transport. In addition to regulating cell elongation further back along the root, auxin also positively regulates levels of the PIN phosphatase subunit RCN1, thereby diminishing the magnitude of AGO. This causes the equilibrium between angle-dependent gravitropic- and angle-independent antigravitropic auxin flux to occur at a more vertical setpoint angle. (I) Tropic response to displacement either above or below GSA is driven by angle-dependent changes in downward gravitropic auxin flux acting in tension with a more constant, angle-independent upward antigravitropic auxin flux. The thickness of red and green arrows signify relative auxin flux.

To understand if auxin regulates lateral root GSA directly via RCN1 levels, we analysed the effect of auxin on the abundance of an RCN1::RCN1:GFP translational reporter by confocal microscopy. We found that treatment with 50 nM IAA for 4 hours significantly increased GFP signal in both lateral and primary root columella cells (Fig. 4C,D; S4D,E). Analysis of *RCN1* transcript levels in lateral root tips treated with 50 nM of IAA over a time course between 2 and 8 hours showed that auxin had no significant effect on *RCN1* expression compared to mock-treated lateral roots (Fig. S4F), indicating that the effect of auxin on RCN1 protein levels is post-transcriptional in nature. These data suggest that auxin is able to regulate lateral root GSA through a signalling pathway that is dependent on RCN1 stabilisation.

Because the expression of RCN1 solely in the columella had the same effect as exogenous auxin treatment on lateral root GSA, we decided to examine the polarity of PIN3:GFP and PIN7:GFP within the columella cells of lateral roots either in the *ARL2::RCN1* background, or treated with 50 nM of IAA or 5F-IAA, an auxin analogue acting specifically through the TIR1 signalling pathway (Robert et al., 2010). Under all of these conditions, we found that the polarity of PIN3:GFP was significantly shifted towards the lower side of lateral root columella cells relative to WT or mock controls (Fig. 4. F,G; Fig. S4G,H). In contrast, the polarity of PIN7:GFP was unaffected both in the *ARL2::RCN1* background and by auxin treatment (Fig. 4. F,G; Fig. S4G,H). These data underline the differences in the subcellular targeting of PIN3 and PIN7 in the gravity-sensing cells. They also indicate that auxin can act to induce more vertical lateral root GSAs by stabilising RCN1 in the columella, thereby reducing the pool of phosphorylated PIN3 and hence the capacity for upward, antigravitropic auxin flux from the lateral root tip. This reduction in upward auxin flux and the angle-dependence of graviresponse in the lateral root means that the equilibrium between gravitropic and antigravitropic auxin fluxes occurs at a smaller angle of displacement from the vertical, producing a steeper GSA.

## Discussion

The ability of plants to maintain their lateral organs at specific GSAs appears to be a complex problem requiring both the monitoring of multiple growth angles and the capacity to reversibly control gravitropic responses both with and against the gravity vector. Here we have shown that non-vertical GSAs in lateral roots arise from the interaction of just two phenomena— angle-dependent gravitropic response and an angle-independent antigravitropic offset— mediated at the level of PIN phosphorylation in the gravity-sensing columella cells (Fig. 4H,I).

The demonstration of quantitative, angle-dependent variation in lateral root gravitropic response is significant because while the angle of growth is set by the magnitude of the AGO, it is the capacity of gravitropic response to increase with displacement from the vertical that provides a means to maintain that angle of growth. The central importance of angle-dependence in the maintenance of non-vertical GSAs contrasts with its apparent dispensability for achieving vertical growth in primary organs. Although angle-dependence contributes to limiting the overshooting in primary roots returning to the vertical following displacement, it is perhaps more likely that the adaptive significance of angle-dependence as a phenomenon lies more in its capacity to sustain gravity-dependent non-vertical growth than to finesse gravitropic response in vertically-growing primary roots.

At the molecular level, several data point to events within the gravity-sensing cells as being critical for GSA maintenance. First, post-stage II lateral roots growing at non-vertical GSAs do not show an asymmetry in auxin distribution between the upper and lower halves of the root (Fig. 1C,D; Fig. S1G). Second, reorientation either above or below GSA induces the generation of a lower- or upper-side-biased auxin asymmetry in the lateral root respectively, including tissues immediately flanking the gravity-sensing cells (Fig. 1C,D; Fig. S1G). Third, these patterns of auxin distribution and redistribution can be traced back to changes in PIN3 and PIN7 polarity within the columella (Fig. S2B-E). Fourth, expression of the PIN phosphatase subunit RCN1, specifically in the columella, is sufficient to alter both the subcellular distribution of PIN3 and lateral root GSA (Fig. 4B,E-G). These observations also emphasize that the maintenance of non-vertical GSAs, including both upward and downward growth, can be understood entirely within the framework of the Cholodny-Went model of tropic growth. Further, because both gravitropic and activities share the same molecular components, it is the relative magnitude of each that determines angle of growth, independent of the overall levels of auxin and PIN proteins.

At the organ-level, the pattern of cell elongation on the upper and lower sides of lateral roots gravistimulated above and below their GSA shows that the steering of growth relative to gravity is driven only by changes on the lower side of the root (Fig. 1E,F). Together with the quantification of auxin distribution in graviresponding lateral roots, these data indicate that the ability to generate upward growth in lateral roots requires a capacity to respond to lower levels of auxin with increased cell expansion in the elongation zone.

The concept of gravitropic and antigravitropic activities acting in tension to generate gravity-dependent non-vertical growth becomes less abstract when thought of in the mechanistic terms of the PIN proteins that mediate auxin efflux from the gravity-sensing columella cells. PIN3 and PIN7 in the columella of stage III lateral roots growing at GSA have distinct polarity patterns that together generate a state of symmetrical upper-vs lower-side auxin distribution. While the majority of lateral roots analysed showed a bipolar distribution of PIN3:GFP (~45%), there was also a significant proportion of roots showing a downward bias (~35%). In contrast, PIN7:GFP showed a conspicuous upward bias (~52%) (Fig. 2A,B,E). This distinctive polarity pattern for PIN7 is reflected in the reorientation kinetics of single and double columella PIN mutants. Lateral roots of *pin3* and *pin3 pin4* mutants display rapid upward-bending relative to downward bending, a phenomenon that is lost in the absence of PIN7 (Fig. 2F-I; Fig. S1J). While these data indicate that PIN3 and PIN7 make distinct contributions in mediating gravitropic and antigravitropic auxin flux from the columella, they are not exclusive for one or the other. For both PIN3:GFP and PIN7:GFP, reorientation either above and below GSA causes shifts in polarity in lateral root gravity-sensing cells that are consistent with the observed changes in auxin distribution in both downward and upward-bending roots (Fig. S2B-E).

The polarity patterns of PIN3 and PIN7 in the lateral root columella were further explored in experiments designed to understand if there are subcellular differences in PIN protein-plasma membrane dynamics, particularly on the upper and lower sides of the cells. This analysis showed that PIN3:GFP and PIN7:GFP signals persisted on membranes of the former upper side of the columella relative to the lower side (Fig. 3A-D). The persistence of an upper side signal is more pronounced for PIN7:GFP compared to PIN3:GFP, possibly reflecting the inherent difference in polarity patterns of the two proteins. The relatively rapid reduction in PIN3:GFP and PIN7:GFP from the former lower membrane of the gravity-sensing cells in flip assays (Fig. 3A-D) is significant because it supports the idea that the loss of gravitropic stimulation, as in the case of a lateral root moved to the vertical, is associated with a rapid reduction in PIN protein abundance and hence, auxin transport capacity from the lower membrane of the statocyte relative to the upper side.

In addition to accounting for differences in the kinetics of change in gravitropic and antigravitropic activities, the disparity in PIN protein dynamics on the upper and lower sides of columella cells also raises the possibility of a parsimonious model of cellular polarity that avoids the requirement to specify ‘up’ and ‘down’ domains within the cell separately. If statolith sedimentation simply defines a ‘down’ domain in each cell, the remainder of the cell can be said to be in a ‘not-down’ state. However, since auxin transport from the lateral plasma membranes of the statocyte are perpendicular to the gravity vector, it is only the relative magnitude of auxin transport from the down and the opposing not-down/up faces of the cells that determine growth trajectory in the vertical plane.

Several molecular and genetic data indicate that the subcellular partitioning of PIN3 to down and not-down regions of columella cells is regulated by the phosphorylation of sites within its cytoplasmic loop, including those targeted by the PID/WAG class of AGC kinases. Mutation of serines 316, 317 and 321 to the phosphomimic aspartic acid results in a upper-side shift in PIN3:GFP signal and concomitant change to a more horizontal GSA, while non-phosphorylatable alanine substitutions result in a shift to a more vertical GSA (Fig. 3E-I). At this point we cannot distinguish between the effect of the serine to alanine mutations at residues 316, 317 and 321 of PIN3 as being to promote targeting to ‘down’ domains of the cell or to inhibit their phosphorylation-dependent targeting to not-down domains. The same applies in the case of the mutation of these residues to aspartic acid, in that the effect could be either or both the active targeting of phosphovariant PIN3:GFP to not-down domains or the inhibition of its recruitment to down domains. Whatever the case, these data point to regulatory events involving these phosphosites as being important for PIN protein targeting in the gravity-sensing cells of the root and hence regulation of GSA.

The idea that phosphoregulation of columella PINs is central to GSA control is further supported by the finding that loss of RCN1 function, a phosphatase subunit involved in PIN dephosphorylation, results in a more horizontal GSA phenotype. Importantly, this phenotype is rescued by expression of *RCN1* solely in the columella, confirming that the RCN1 activity relevant to GSA control occurs in the gravity-sensing cells (Fig. 4E). The overexpression of RCN1 in the columella, in a wild-type background, induces a more vertical GSA and a concomitant downward shift in PIN3:GFP but strikingly, not in PIN7:GFP, again highlighting the differences in the subcellular targeting of these proteins (Fig. 4F,G).

Our data also show that RCN1 activity in the columella is part of the mechanism by which auxin induces more vertical GSA. *rcn1* lateral roots are almost entirely resistant to the effect of auxin on GSA and auxin treatment increases RCN1 levels in the columella, inducing a downward shift in PIN3:GFP, but not PIN7:GFP, consistent with the effects of *RCN1* overexpression. Together, these data provide compelling support for the idea that control of RCN1 levels and hence PIN3 polarity in the columella is central to auxin’s ability to regulate lateral root GSA.

## Conclusions

Our work has shown how the patterns of growth angle control observed in lateral roots arise from the interaction of angle-dependent gravitropic response and angle-independent, auxin-repressible antigravitropic offset, both mediated at the level of lateral root columella PIN proteins. The auxin-tunable nature of the AGO provides a means for the developmental and environmental modulation of angled growth in lateral organs (Roychoudhry et al., 2017). At this point the molecular basis of the cell biological differences between PIN3 and PIN7 in lateral root statocytes is not clear. The fact that the subcellular distribution of PIN7 in the columella is unaffected by RCN1 activity suggests that the reason for the distinct upper side bias of PIN7 is either not related to phosphorylation or at the very least, involves phosphorylation of sites that are not subject to regulation by RCN1. For PIN3, although phosphorylation of serines 316, 317 and 321 is functionally relevant to the control of its polarity in the columella, we do not know if other phosphosites in its cytoplasmic loop, which are targeted by RCN1, contribute to GSA control. We also do not exclude other mechanisms contributing to GSA control in the lateral root. In particular, it will be interesting to establish the level at which LAZY proteins, recently shown to play a role in primary and lateral root gravitropism (Ge & Chen, 2016; Guseman et al., 2017; Taniguchi et al., 2017; Yoshihara & Spalding; 2017), input into the control of PIN polarity in lateral root statocytes and how their activity integrates with the phosphoregulation of PIN3 identified here. Perhaps the most conspicuous open question at hand is a very old one, that of how statolith sedimentation is turned into asymmetry in PIN localisation and activity within statocytes. In the context of GSA control, the question relates to understanding the basis of angle-dependent graviresponse that is so crucial to the maintenance of non-vertical GSAs. By highlighting the evolutionary and adaptive significance of such phenomena, the model of GSA control proposed here provides not only practical tools but also fresh approaches to tackling these fascinating and important questions in plant biology.

## Materials and methods

### Plant materials

All Arabidopsis seed stocks are in the Col-0 background unless otherwise stated. R2D2 (Liao et al, 2015) is in the Utrecht background. The *pin3-3, pin7-2, pin3-5 pin7-1 [pin3 pin4], pin3-5 pin4-3 [pin3 pin4], pin3-5 pin4-3 pin7-1 [pin3 pin4 pin7]* (Friml et al., 2002; Friml et al., 2003), PIN3:GFP, PIN7:GFP (Kleine-Vehn et al., 2010), *rcn1* (Rashotte et al., 2001), *wag1 wag2* (Santner & Watson, 2006), PIN3:YFP, PIN3S>A:YFP and PIN3S>D:YFP (Grones et al., 2018), *RCN1::RCN1:GFP [PP2AA::PP2AA:GFP], PID::PID:Venus, WAG1::GUS, WAG2::GUS* (Dhonushke et al., 2010), PIN3:S4S5A:YFP [and PIN3:YFP control] (Zourelidou et al., 2014), DII-Venus (Band et al., 2012), *TIR1::TIR1:Venus, AFB2::AFB2:Venus*, and *AFB3::AFB3:Venus* (Roychoudhry et al., 2017) lines have been described previously. Ws-0 seeds were obtained from the Nottingham Arabidopsis Stock Centre. The *ARL2:RCN1* construct was generated by cloning 2.5 kb of the *ARL2* promoter upstream of the *RCN1* coding sequence using a multiplex gateway cloning strategy (Invitrogen) into a pALLIGATOR V destination vector. The *ARL2::GFP* construct was generated by cloning 2.5 kB of the *ARL2* promoter sequence into a modified pGreen 0229 vector containing GFP cloned upstream of a NOS terminator. Transformation of these constructs to Arabidopsis was accomplished via *Agrobacterium tumefaciens* (strain GV3101)-mediated infiltration by floral dip. *ARL2::RCN1* PIN3:GFP and *ARL2:RCN1* PIN7:GFP lines were generated by crossing.

### Reorientation experiments

12-day-old seedlings grown vertically on 120 mm square ATS media plates under light and temperature regimes described above were reorientated by appropriate angles in darkness. Images were captured automatically at described intervals using a Canon 700D digital camera and infra red illumination using the ‘Image Capture’ software in OS El Capitan on a 2013 MacBook Pro in order to nullify any phototropic effects. Root tip angles were quantified using ImageJ.

### Maintenance of constant gravistimulation and measurement of root orientation

Roots were illuminated with an infrared light-emitting diode (Radio Shack, Fort Worth, TX) and imaged with a CCD camera interfaced to a computer via a frame grabber card (Imagenation Corp., Beaverton, OR). A computer feedback system connected to a rotary stage (Mullen et al., 2000) was used to measure the orientation of the root apex and constrain it to that initial orientation prior to gravistimulation by making corrections every 45 s. Following reorientation, the root tip was constrained at the new orientation to maintain a constant gravistimulus throughout the experiment. Gravitropic curvature was measured as the rotation of the stage necessary to maintain the root tip at a constant orientation.

### qRT-PCR for RCN1 expression

RNA was extracted from the lateral and primary root tips of 12 day-old wild-type Col-0 plants grown on ATS media overlaid with Sefar Nitex mesh using the Qiagen RNAeasy kit according to the manufacturer’s instructions. cDNA was synthesized from the isolated RNA using oligo dT primers and Superscript II reverse transcriptase (Invitrogen). qPCR was performed using the Bio-Rad CFX Connect Real-Time System (Bio-Rad). GAPDH was used as an internal control.

### Analysis of lateral root GSA

Lateral root GSA was analysed as previously described in Roychoudhry et al., 2017.

### Quantification of bending during clinorotation

12 day old Arabidopsis seedlings growing on 9 cm petri dishes were mounted on a horizontal clinostat as described in Roychoudhry & Kepinski, 2015. The plates were photographed at hourly time intervals using a Canon Powershot G90 digital camera.Root tip angles were quantified using ImageJ.

### Confocal microscopy

10-12 day old marker seedlings grown on ATS or half MS media in standard tissue culture conditions (20-22 °C 16h day, 8h dark) were imaged at 20X resolution with the 480 nm and 540 nm lasers using a Zeiss LSM 710 inverted confocal microscope. For vertical stage confocal microscopy and gravistimulation, the imaging setup described in Von Wangenheim et al., (2017) was used. All laser power and gain settings were consistent across images. Briefly, PIN:GFP markers were imaged using a series of Z stacks and fluorescence intensity across external membranes was quantified using ImageJ as described in Grones et al., 2018. Each experiment was performed three times with at least 6 roots for each experiment. For cell length quantification, wild-type Col-0 plants on ATS grown on ATS media were reorientated for a period of 6 hours. The entire root system was mounted on a glass slide and counter stained with propidium iodide prior to imaging stage III lateral roots. Epidermal atrichoblast cell length was quantified using ImageJ. The experiment was performed three times with at least 6 roots at each orientation per experiment. For DII-Venus and R2D2, excluding the lateral root cap, nuclear fluorescence was measured in ten consecutive epidermal cells within the two outermost flanking cell files, beginning from the root tip for each root. Experiments were performed three times with at least ten root tips for each orientation per experiment. For R2D2, nuclear fluorescence intensity was measured across both GFP and mTOMATO channels. For each nucleus, the ratio of GFP/mTOMATO signal was determined. Geometric means and standard errors of the ratios were calculated for both young, and older lateral roots. Student’s T-test was performed to evaluate statistical differences between the geometric means of the data obtained.

### Primer sequences

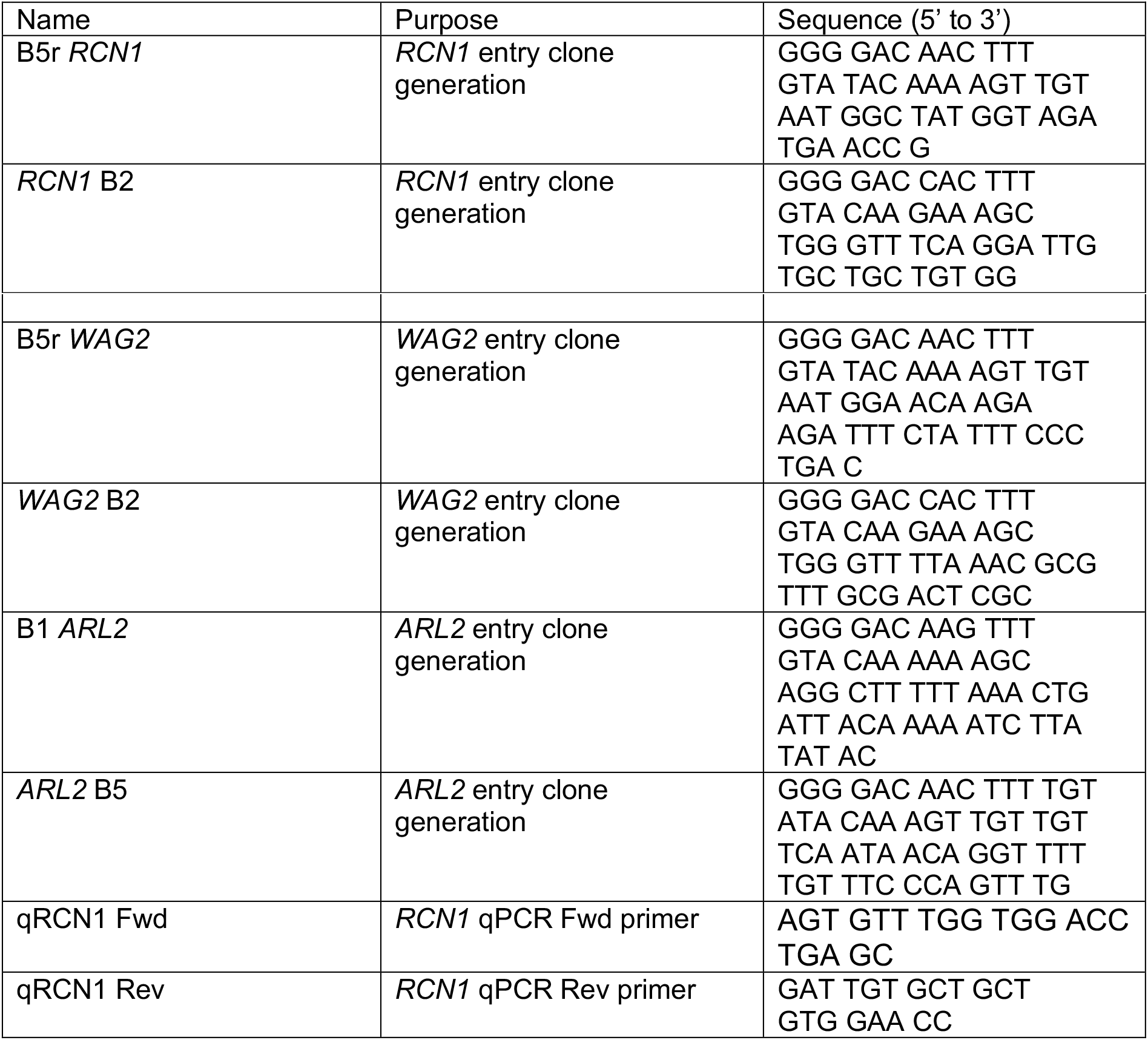

## Acknowledgments

We thank Dolf Weijers, Claus Schwechheimer, and Remko Offringa for generous sharing of published and unpublished materials and we are grateful to Marta Del Bianco and Ottoline Leyser for critical reading of the manuscript. This work was supported by the BBSRC (grants BB/N010124/1 and BB/R000859/1 to SK).

## Author contributions

S.R performed the majority of experiments and analysed the data except: K.S-F generated and analysed the R2D2 data. H.L.G analysed GSA phenotypes of the *wag1 wag2* mutants. P.G and J.F generated the PIN3:YFP phosphovariant lines. C.W, J.M and R.H generated the lateral root angle-dependence data. J.F provided critical experimental suggestions and feedback on the draft manuscript. S.R. and S.K wrote the manuscript. All authors commented on and approved the manuscript.

## References

Adamowski, M. & Friml, J. PIN-dependent auxin transport: action, regulation, and evolution Plant Cell 27, 20–32 (2015).

Band, L.R., Wells, D.M., Larrieu, A., Sun, J., Middleton, A. M., French, A. P., et al. Root gravitropism is regulated by a transient lateral auxin gradient controlled by a tipping-point mechanism. Proc. Natl. Acad. Sci. USA 109, 4668–73 (2012).

Baster, P., Robert, S., Kleine-Vehn, J., Vanneste, S., Kania, U., Grunewald, W., De Rybel, B., Beeckman, T. & Friml, J. SCF(TIR1/AFB)-auxin signalling regulates PIN vacuolar trafficking and auxin fluxes during root gravitropism. EMBO J. 32, 260–274 (2013).

Blancaflor, E. & Masson, P. Plant Gravitropism. Unravelling the ups and downs of a complex process. Plant Physiol. 133, 1677–90 (2003).

Blilou, I., Xu, J., Wildwater, M., Willemsen, V., Paponov, I., Friml, J., Heidstra, R., Aida, M., Palme, K. & Scheres, B. The PIN auxin efflux facilitator network controls growth and patterning in Arabidopsis roots. Nature 433, 39–44 (2005).

Dhonukshe, P., Huang, F., Galvan-Ampudia, C. S., Mahonen, A. P., Kleine-Vehn, J., Xu, J., Quint, A., Prasad, K., Friml, J., Scheres, B. & Offringa R. Plasma membrane-bound AGC3 kinases phosphorylate PIN auxin carriers at TPRXS(N/S) motifs to direct apical PIN recycling. Development 137, 3245–55 (2010).

Digby, R.D. & Firn, J. The gravitropic set-point angle (GSA): the identification of an important developmentally controlled variable governing plant architecture. Plant Cell Environ. 18, 1434–40 (1995).

Fendrych, M., Akhmanova, M., Merrin, J., Glanc, M., Hagihara, S., Takahashi, K., Uchida, N., Torii, K.U. & Friml, J. Rapid and Reversible Root Growth Inhibition by TIR1 Auxin Signalling. Nat. Plants 2018, 4, 453–459.

Feraru, E. & Friml, J. PIN polar targeting. Plant Physiol. 147, 1553–1559 (2008).

Friml, J., Wisniewska, J., Benková, E., Mendgen, K. & Palme, K. Lateral relocation of auxin efflux regulator AtPIN3 mediates tropism in Arabidopsis. Nature, 415, 806–9 (2002).

Friml, J., Yang, X., Michniewicz, M., Weijers, D., Quint, A., Tietz, O., Benjamins, R., Ouwerkerk, P. B.F., Ljung, K., Sandberg, S., Hooykaas, P.J.J., Palme, K. & Offringa, R. (2004). A PINOID-dependent binary switch in apical-basal polar PIN targeting directs auxin efflux. Science 306: 862–865.

Friml J., Vieten A., Sauer M., Weijers D., Schwarz H., Hamann T., Offringa R. & Jürgens G. Efflux-ependent auxin gradients establish the apical–basal axis of Arabidopsis. Nature 426, 147–153 (2003).

Ganguly, A., Sasayama, D. & Cho H.-T. Regulation of the polarity of protein trafficking by phosphorylation. Mol. Cells 33, 423–30 (2012).

Geldner N., Denervaud-Tendon V., Hyman D.L., Mayer, U., Stierhof Y.D. & Chory J. Rapid, combinatorial analysis of membrane compartments in intact plants with a multicolor marker set. Plant J. 59, 169–178 (2009).

Ge L. & Chen R. Negative gravitropism in plant roots. Nature Plants 2, 16155–59 (2016).

Glanc M., Fendrych M. & Friml J. Mechanistic framework for cell-intrinsic re-establishment of PIN2 polarity after cell division. Nature Plants 4, 1082–1092 (2018).

Grones P., Abas M., Hajny J., Jones A., Weidmann S., Kleine-Vehn J. & Friml J. PID/WAG-mediated phosphorylation of the Arabidopsis PIN3 auxin transporter mediates polarity switches during gravitropism. Sci. Rep 8, 10279 (2018).

Guseman J.M., Webb K., Srinivasan C. & Dardick C. DRO1 influences root system architecture in Arabidopsis and *Prunus* species. Plant J. 89, 1093–1105 (2017).

Guyomarc’h S., Leran S., Auzon-Carpe M., Perrine-Walker F., Lucas M. & Laplaze L. Early development and gravitropic response of lateral roots in Arabidopsis thaliana. Phil. Trans. R. Soc. Lond. B. Biol. Sci. 367, 1509–1516 (2012).

Haberlandt G. Ueber die perzeption des geotropischen reizes. Ber Dtsch Bot Ges 18, 261–272 (1900).

Harrison, B.R. & Masson, P.H. ARL2, ARG1 and PIN3 define a gravity signal transduction pathway in root statocytes. Plant J. 53, 380–392 (2008).

Hou G., Mohamalawari D.R. & Blancaflor E.B. Enhanced gravitropism of roots with a disrupted cap actin cytoskeleton. Plant Physiol. 131, 1360–1373 (2003).

Hou G., Kramer V.L., Wang Y., Chen R., Perbal G., Gilroy S. & Blancaflor E.B. The promotion of gravitropism in *Arabidopsis* roots upon actin disruption is coupled with the extended alkalinization of the columella cytoplasm and a persistent lateral auxin gradient. Plant J. 39, 113–125 (2004).

Kiss J.Z., Miller K.M., Ogden L.A. & Roth K.K. Phototropism and gravitropism in lateral roots of Arabidopsis. Plant Cell Physiol. 43, 35–43 (2002).

Kleine-Vehn, J., Ding, Z., Jones, A.R., Tasaka, M., Morita, M.T. & Friml, J. Gravity-induced PIN transcytosis for polarization of auxin fluxes in gravity-sensing root cells. Proc Natl Acad Sci. USA 107, 22344–22349 (2010).

Liao, C.Y., Smet, W., Brunoud, G., Yoshida, S., Vernoux, T. & Weijers, D. Reporters for sensitive and quantitative measurement of auxin response. Nat. Methods 12, 207–210 (2015).

Löfke, C., Scheuring, D., Dünser, K., Schöller, M., Luschnig, C. & Kleine-Vehn, J. Tricho- and atrichoblast cell files show distinct PIN2 auxin efflux carrier exploitations and are jointly required for defined auxin-dependent root organ growth. J. Exp. Bot. 66, 5103–5112 (2015).

Michniewicz M., Zago M. K., Abas L., Weijers D., Schweighofer A., Meskiene I., Heisler M. G., Ohno C., Zhang J., Huang F., Schwab R., Weigel D., Meyerowitz E. M., Luschnig C., Offringa R. & Friml, J. Antagonistic regulation of PIN phosphorylation by PP2A and PINOID directs auxin flux. Cell 130, 1044–1056 (2007).

Mullen J.P. & Hangarter R. Genetic analysis of the gravitropic set-pointangle in lateral roots of Arabidopsis. Adv. Space Res. 31, 2229–22364 (2003).

Mullen J., Wolverton C., Ishikawa H. & Evans M.L. Kinetics of Constant Gravitropic Stimulus Responses in Arabidopsis Roots Using a Feedback System. Plant Physiol. 23, 665–670 (2000).

Nemec, B. Ueber die art der wahrnehmung des schwek-raftreizes bei den pflanzen. Ber Dtsch Bot Ges 18, 241–245 (1900).

Ottenschläger I., Wolff P., Wolverton C., Bhalerao R.P., Sandberg G., Ishikawa H., Evans M. & Palme, K. Gravity-regulated differential auxin transport from columella to lateral root cap cells. Proc. Natl Acad. Sci. USA 100, 2987–2991 (2003).

Paciorek, T., Zazímalová, E., Ruthardt, N., Petrásek, J., Stierhof, Y.D., Kleine-Vehn, J., Morris, D.A., Emans, N., Jürgens, G., Geldner, N. & Friml, J. Auxin inhibits endocytosis and promotes its own efflux from cells. Nature 435, 1251–1256 (2005).

Pernisova M., Prat T., Grones P., Harustiakova D., Matonohova M., Spichal L., Nodzynski T., Friml, J. & Hejatko J. Cytokinins influence root gravitropism via differential regulation of auxin transporter expression and localization in Arabidopsis. New Phytol. 212, 497–509 (2016).

Rakusova H., Gallego-Bartoleme J., Vanstraelen M., Boisivon H., Alabadi D., Blazquez M.A., Benkova E., Friml J. Polarisation of PIN3-dependent auxin transport for hypocotyl gravitropic response in Arabidopsis thaliana. Plant J. 67, 817–826 (2011).

Rashotte, A. R., Brady, S. R., Reed, R. C., Ante, S. J. & Muday, G. K. Basipetal auxin transport is required for gravitropism in roots of Arabidopsis. Plant Physiol. 122, 481–490 (2000).

Rehman, A., Takahashi, M., Shibasaki, K., Wu, S., Inaba, T., Tsurumi, S. & Baskin, T. Gravitropism of Arabidopsis thaliana roots requires the polarization of PIN2 toward the root tip in meristematic cortical cells. Plant Cell 22, 1762–76 (2010).

Robert, S., Kleine-Vehn J., Barbez E., Sauer M., Paciorek T., Baster P., Vanneste S., Zhang J., Simon, S., Covanova, M., Hayashi, K., Dhonukshe P., Yang Z., Bednarek S.B., Jones A.M., Luschnig C., Aniento F., Zazimalova E. & Friml J. ABP1 Mediates Auxin Inhibition of Clathrin-Dependent Endocytosis in Arabidopsis. Cell 143, 111–121(2010).

Roychoudhry, S., Del Bianco M., Kieffer, M. &Kepinski, S. (2013). Auxin controls gravitropic setpoint angle in higher plant lateral organs. Curr. Biol. 23, 1497–1504 (2013).

Roychoudhry, S. &Kepinski, S. Root and shoot branch growth angle control - The wonderfulness of lateralness. Curr. Op. Plant Biol. 23, 124–131 (2015).

Roychoudhry, S. &Kepinski, S. The analysis of gravitropic setpoint angle control in Arabidopsis. Methods in Mol. Biol. 1309:31–41 (2015).

Roychoudhry, S., Kieffer, M., Del Bianco, M., Liao, C.Y., Weijers, D. &Kepinski, S. The developmental and environmental regulation of gravitropic setpoint angle in Arabidopsis and bean. Sci. Rep. 7, 42664 | DOI: 10.1038/srep42664 (2017)

Ruiz Rosquete, M., von Wangenheim, D., Marhavy, P., Barbez, E., Stelzer, E.H.K., Benkova, E., Maizel, A. & Kleine-Vehn, J. An auxin transport mechanism restricts positive orthogravitropism in lateral roots. Curr. Biol. 23, 817–822 (2013).

Sachs, J. Über orthotrope und plagiotrope Pflanzenteile. Arb Bot Inst Würzburg 2, 226–284 (1882).

Sack, F.D. & Kiss, J.Z. Rootcap structure in wild-type and in a starchless mutant of Arabidopsis. Am. J. Bot. 76, 454–64 (1989).

Santner, A.A. & Watson, J.C. The WAG1 and WAG2 protein kinases negatively regulate root waving in Arabidopsis. Plant J. 45, 752–764 (2006).

Sukumar, P., Edwards, K. F., Rahman, K., DeLong, A. & Muday, G.K. PINOID Kinase Regulates Root Gravitropism through Modulation of PIN2-Dependent Basipetal Auxin Transport in Arabidopsis. Plant Physiol. 150, 722–735 (2009).

Taniguchi, M., Furutani, M., Nishimura, T., Nakamura, M., Fushita, T., Iijima, K., Baba, K., Tanaka, H., Toyota, M., Tasaka, M. & Morita, M.T. The Arabidopsis LAZY1 family plays a key role in gravity signaling within statocytes and in branch angle control of roots and shoots. Plant Cell 29, 1984–1999 (2017).

Wisniewska, J., Xu, J., Seifertova, D., Brewer, P. B., Ruzicka, K., Blilou, I., Rouquie, D., Benkova, B., Scheres, B. & Friml J. Polar PIN localization directs auxin flow in plants. Science 312, 883–890 (2006).

Vieten, A., Vanneste, S., Wisniewska, J., Benkova, E., Benjamins, R., Beeckman, T., Luschnig, C. &Friml, J. Functional redundancy of PIN proteins is accompanied by auxin-dependent cross-regulation of PIN expression. Development 132, 4521–4531 (2006).

von Wangenheim, D., Hauschild, R., Fendrych, M., Barone, V., Benkova, E. & Friml J. Live tracking of moving samples in confocal microscopy for vertically grown roots. eLife 6, e26792 (2017).

Yoshihara, T., Spalding, E.P., Iino M. AtLAZY1 is a signaling component required for gravitropism of the Arabidopsis thaliana inflorescence. Plant J. 74, 267–279 (2013).

Yoshihara, T. & Spalding, E.P. (2017) LAZY Genes Mediate the Effects of Gravity on Auxin Gradients and Plant Architecture. Plant Physiol. 175, 959–969 (2017).

Zhang, J., Nodzyński, T., Pĕnčík, A., Rolčík, J. & Friml, J. PIN phosphorylation is sufficient to mediate pin polarity and direct auxin transport. Proc Natl Acad Sci. USA 107, 918–922 (2009).

Zourelidou, M., Absmanner, B., Weller, B., Barbosa, I. C., Willige, B. C., Fastner, A., Streit, V., Port, S. A., Colcombet, J., de la Fuente van Bentem, S., Hirt, H., Kuster, B., Schulze, W. X., Hammes, U. Z. &Schwechheimer, C. (2014). Auxin efflux by PIN-FORMED proteins is activated by two different protein kinases, D6 PROTEIN KINASE and PINOID. eLife 3 e02860.

