## Supplementary Information for "Antagonistic and auxin-dependent phosphoregulation of columella PIN proteins controls lateral root gravitropic setpoint angle in Arabidopsis"

Roychoudhry et al., Supp. Fig. S1

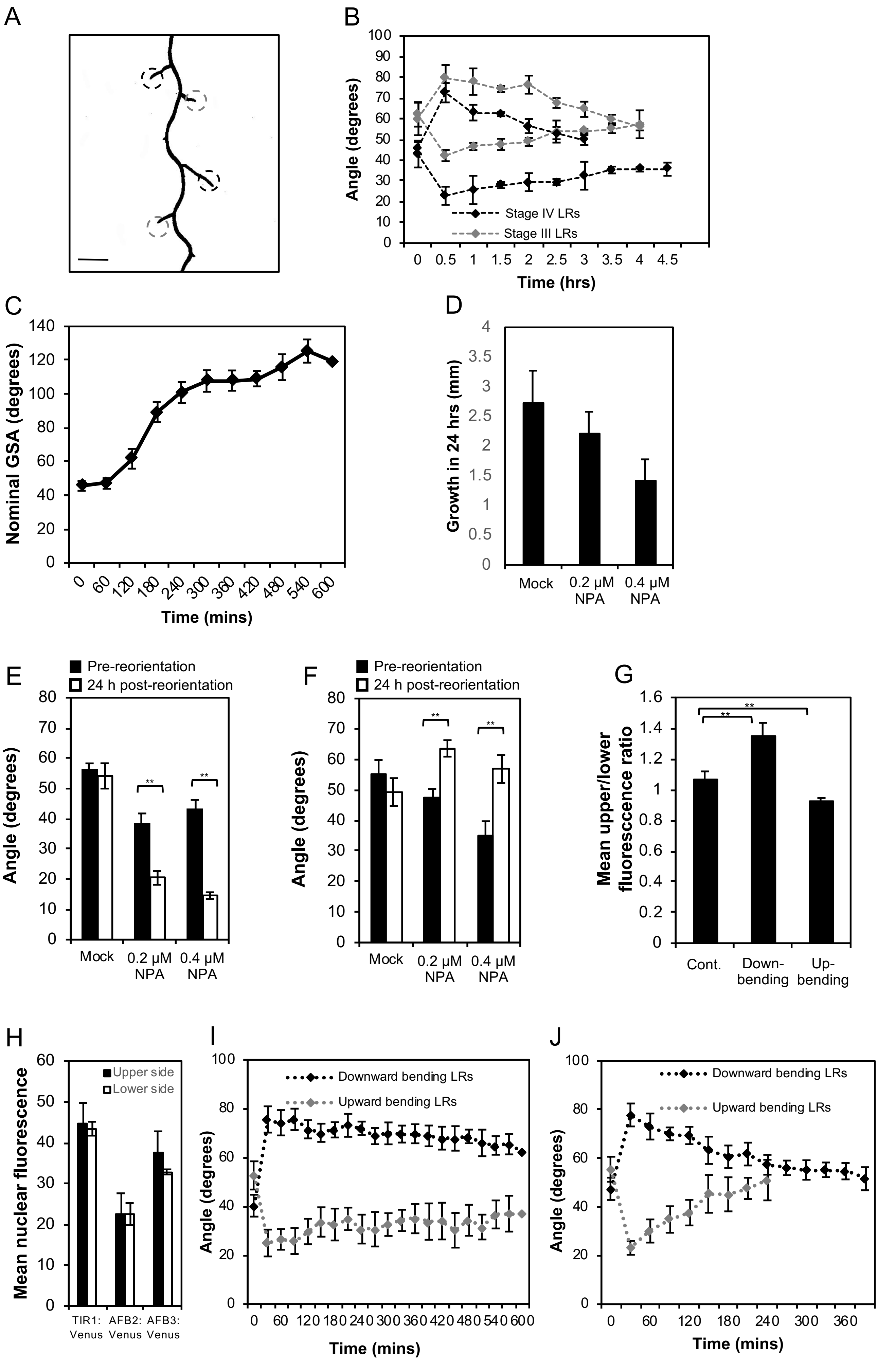

**Figure S1**: (A) Representative 12-day-old Arabidopsis lateral root system showing Stage III (grey dashed circles) and Stage IV (black dashed circles) lateral roots. Scale bar = 5 mm. (B) Reorientation kinetics of Stage III and IV LRs gravistimulated by 30° both above and below their GSA. (C) Nominal GSA of stage III lateral roots at hourly intervals after clinorotation (n = 12) (D) Change in length of stage III lateral roots during mock and NPA treatments. The growth rates of NPA treated roots are not significantly different from mock treated roots (p = 0.0838). Bars represent standard errors of the mean of 3 independent experiments with at least 6 lateral roots in each experiment. (E,F) Mean GSA of mock and NPA treated stage III lateral roots growing at GSA (white bars) and 24 hrs after reorientation by 30° (black bars). Treatment with 0.2 μM and 0.4 μM NPA inhibits lateral root reorientation in both upward (E) and downward (F) directions. (G) Ratio of nuclear DII-Venus signal across upper and lower epidermal cells in lateral roots at GSA (control) and reorientated upwards and downwards. Bars represent standard error of mean nuclear fluorescence. (F) There is no significant difference in mean nuclear fluorescence of TIR1/AFB:Venus in atrichoblast cells on the upper and lower side of stage III lateral roots (p= 0.588, 0.928 and 0.078 respectively). (G) Reorientation kinetics of 12 day old lateral roots in the *pin3 pin4 pin7* triple mutant. Both upward and downward reorientation kinetics are severely delayed in the triple mutant background. (H) Reorientation kinetics of 12 day old lateral roots in the *pin3 pin4* double mutant. PIN7 is required for fast upward bending.

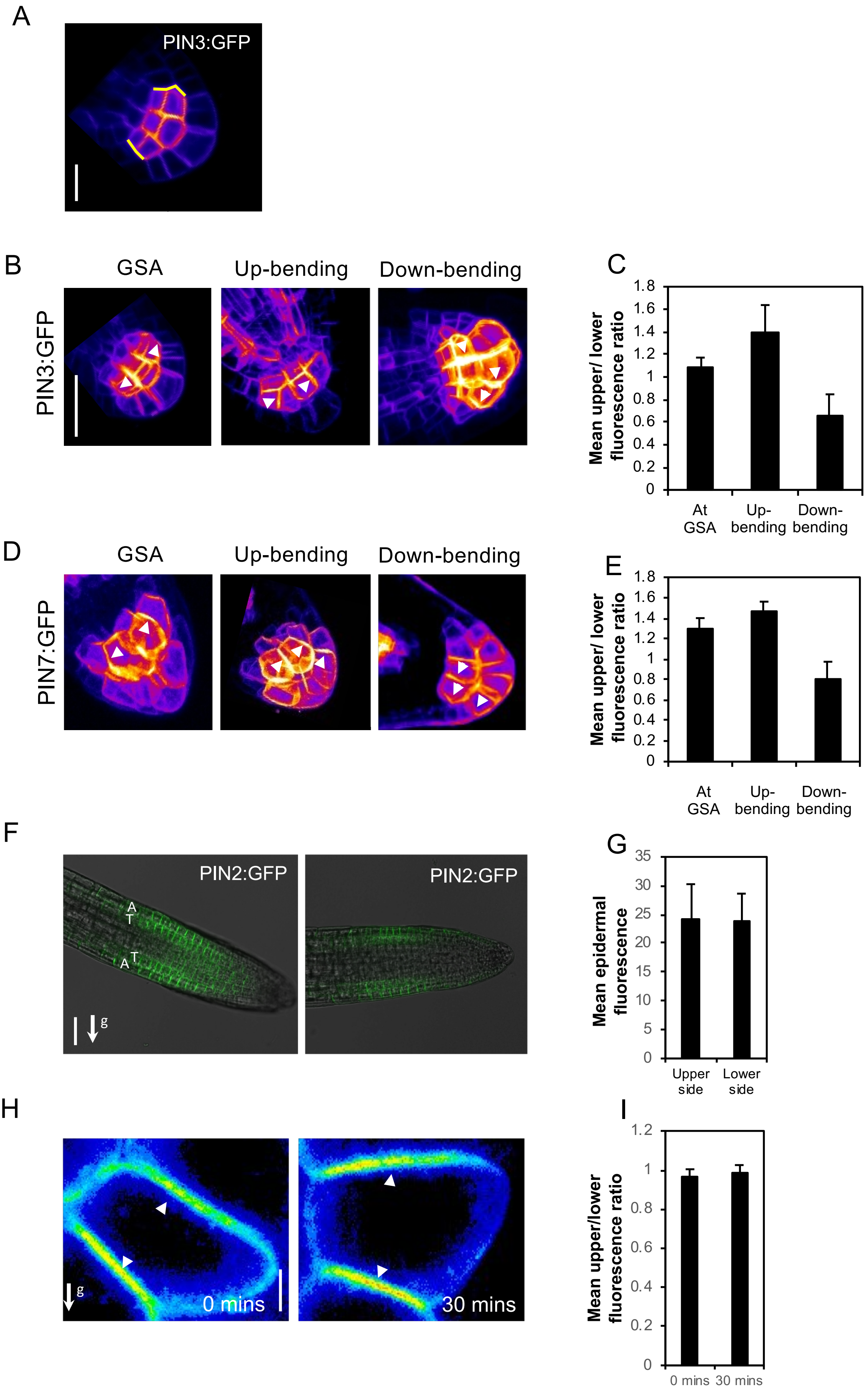


Roychoudhry et al., Supp. Fig. S2

**Figure S2**: (A) Outer membranes (yellow lines) of upper and lower cells of the central columella used for quantification of PIN polarity in a single stack of a PIN3:GFP lateral root. Scale bar = 5 μm (B-E) PIN polarity changes in 12-day old lateral roots reorientated in upward and downward directions. (B,D) PIN3:GFP and PIN7:GFP are predominantly dipolar (PIN3) and apolar (PIN7) in lateral roots at their GSA (right panels). In lateral roots reorientated below their GSA, both PIN3 and PIN7 polarise towards the upper side of columella cells (B,D middle panels) whereas in roots displaced above their GSA, both PINs polarise towards the lower side of columella cells (B,D left panels). White arrowheads indicate polar localisation of PIN:GFP signal. Scale bar = 5 μm. (C,E) Quantification of upper/lower mean GFP signal across external columella cell membranes in lateral roots at GSA and reorientated in both directions. Bars represent standard error of the means. (F) PIN2:GFP signal in lateral roots at their GSA. PIN2:GFP was differentially expressed across trichoblast (T) and atrichoblast (A) cell files (left panel), but not across upper and lower sides of lateral roots (right panel). Scale bar = 20 μm. (G) Quantification of PIN2:GFP in trichoblasts across upper and lower epidermal cell files. Bars represent standard error of the means. (H) The WAVE:YFP plasma membrane marker remains apolar in columella cells at their GSA (0 mins) or 30 mins after 'flipping'. Scale bar = 5 μm (I) Ratio of WAV:YFP plasma membrane polarity in lateral root columella cells at GSA and 30 mins post 'flipping'.

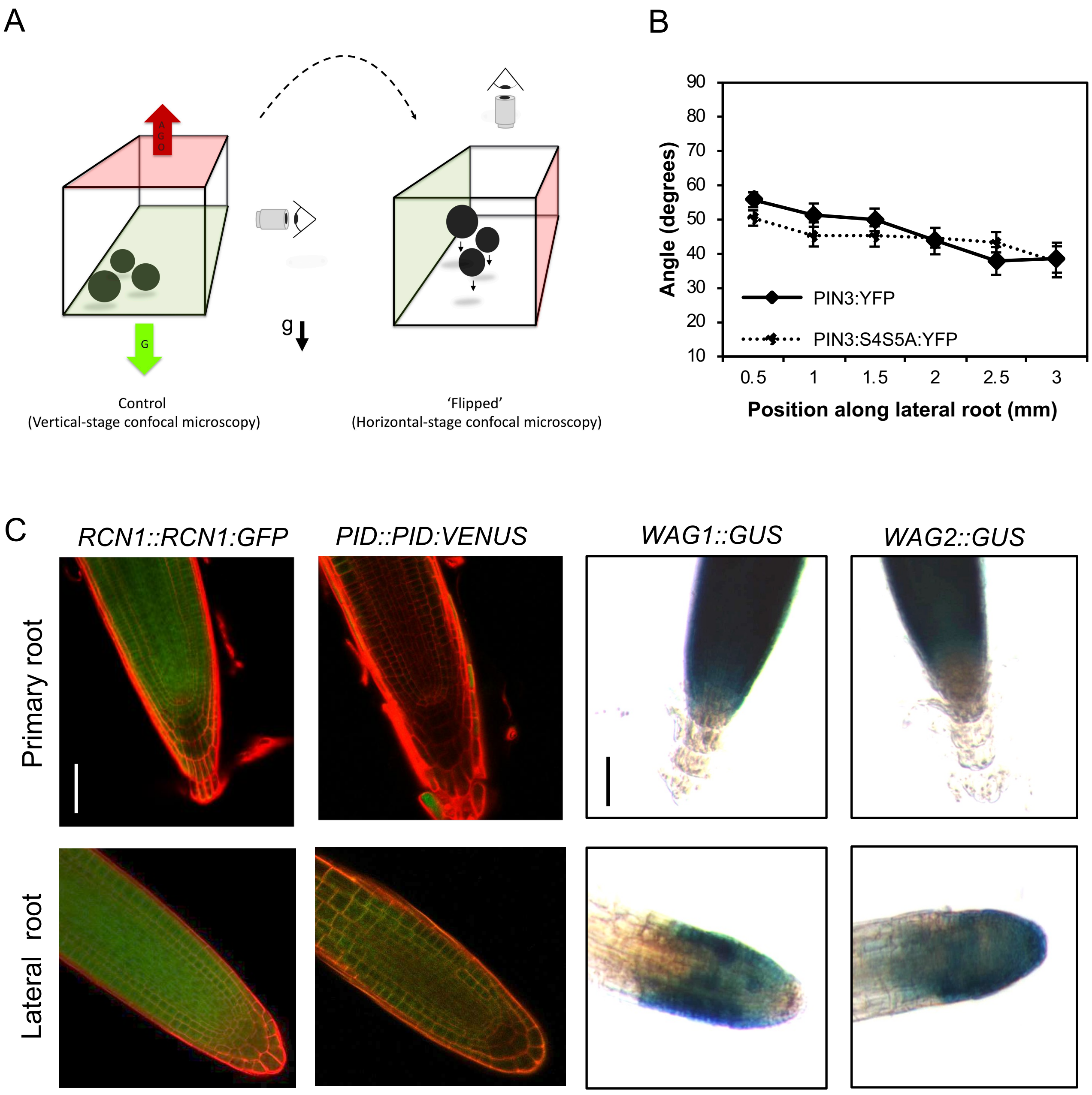


**Figure S3**: (A) Schematic representation of the 'flip' assay designed to study PIN membrane retention kinetics using vertical stage microscopy. (B) Quantification of lateral root GSA in the 12-day-old seedlings of PIN3:YFP D6PK phosphovariant line (PIN3:S4S5:YFP). No significant differences were observed as compared to the PIN3:YFP control. (C) Expression of PP2AA/RCN1 phosphatase subunit and the PID/WAG kinase family in primary and lateral roots. *RCN1::RCN1:GFP* is expressed in both, the primary and lateral root columella cells (left upper and lower panels). In contrast *PID::PID:VENUS* and *WAG1::GUS* are not expressed in the primary or lateral root columella (Centre upper and lower panels). *WAG2::GUS* is absent from the primary root columella, but strongly expressed in the lateral root columella. Scale bar = 30 $\mu$m.

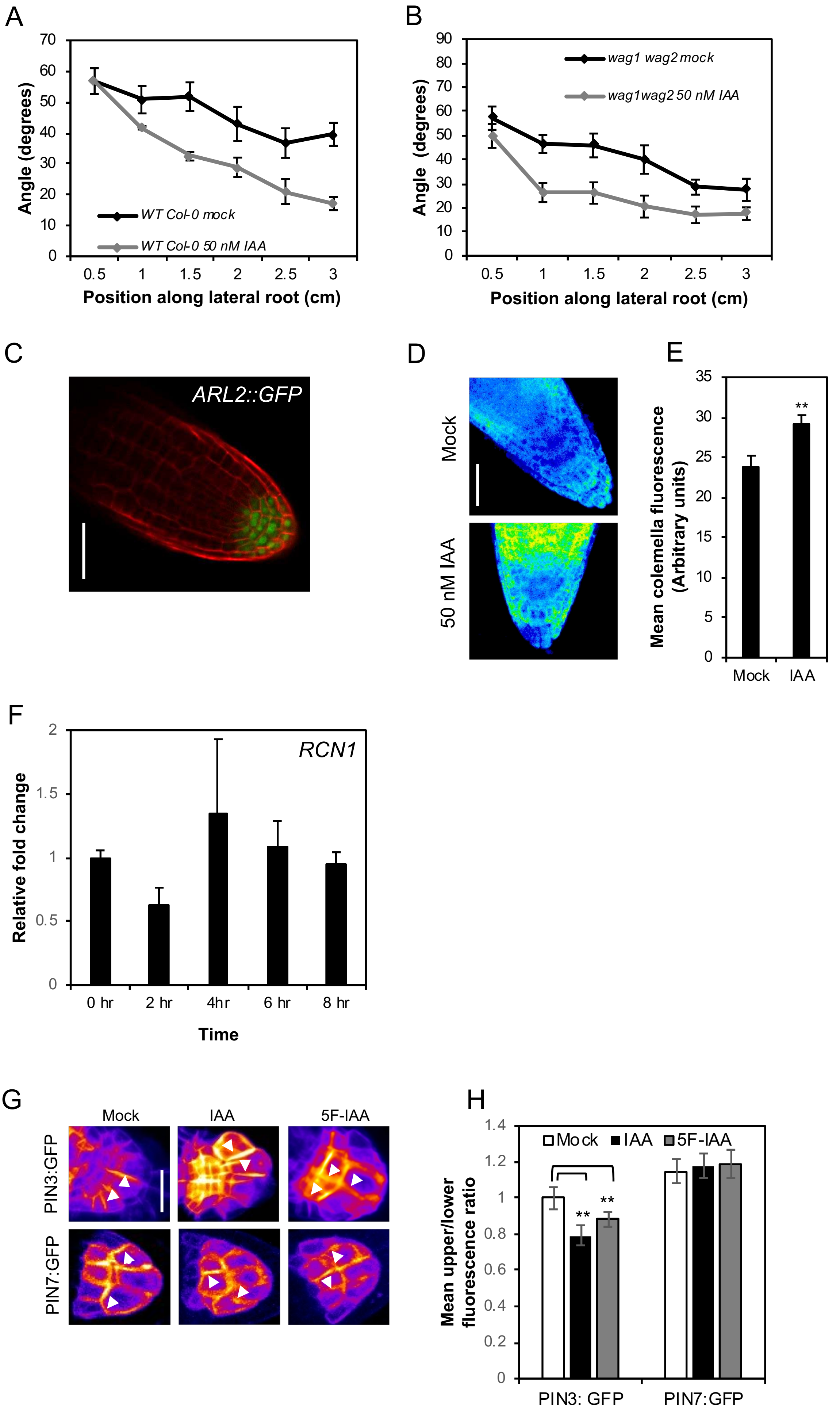


Roychoudhry et al., Supp. Fig. S4

**Figure S4**: (A,B) Effect of auxin treatment on WT Col-0 (A) and *wag1 wag2* (B) Mutant lateral roots. *wag1wag2* lateral roots adopt a more vertical orientation upon auxin treatment. (C) Expression of ARL2::GFP in lateral root columella cells. Scale bar = 50 μm (D,E) Effect of auxin treatment for 4 hours on *RCN1::RCN1:GFP* (*PP2AA::PP2AA:GFP* translational reporter line) primary root columella cells. Auxin treatment leads to a significant increase in RCN1:GFP signal levels (p= 0.007). Scale bar = 20 μm. (F) Effect of 50 nM IAA on *RCN1* transcript levels in lateral root columella cells. No significant increase in *RCN1* levels occurred over an 8 hour time course. Data represent averages from 3 independent experiments with 7-8 root tips harvested for each time point per experiment. Bars represent standard error of the means. (G,H) Treatment with 50 nM IAA or 5F-IAA results in a shift in PIN3:GFP polarity towards the lower side of the columella cell, but has no effect on PIN7:GFP polarity. ** represent a p value of <0.01. Scale bar = 15 μm.
